# Nanobodies as allosteric modulators of Parkinson’s disease-associated LRRK2

**DOI:** 10.1101/2021.08.30.458082

**Authors:** Ranjan K. Singh, Ahmed Soliman, Giambattista Guaitoli, Eliza Störmer, Felix von Zweydorf, Thomas Dal Maso, Laura Van Rillaer, Sven H. Schmidt, Deep Chatterjee, Els Pardon, Stefan Knapp, Eileen J. Kennedy, Jan Steyaert, Friedrich W. Herberg, Arjan Kortholt, Christian J. Gloeckner, Wim Versées

**Affiliations:** VIB-VUB Center for Structural Biology, Pleinlaan 2, 1050 Brussels, Belgium; Structural Biology Brussels, Vrije Universiteit Brussel, Pleinlaan 2, 1050 Brussels, Belgium; Department of Cell Biochemistry, University of Groningen, Groningen, The Netherlands; German Center for Neurodegenerative Diseases (DZNE), Otfried-Müller-Str. 23, D-72076 Tübingen, Germany; Department of Biochemistry, Institute for Biology, University of Kassel, 34132, Kassel, Germany; Institute of Pharmaceutical Chemistry, Goethe-University Frankfurt, Frankfurt, Germany; Structural Genomics Consortium, Buchmann Institute for Molecular Life Sciences (BMLS), Goethe-University Frankfurt, Frankfurt, Germany; Department of Pharmaceutical and Biomedical Sciences, College of Pharmacy, University of Georgia, Athens, Georgia 30602, United States; Core Facility for Medical Bioanalytics, Center for Ophthalmology, Institute for Ophthalmic Research, University of Tübingen, 72076 Tübingen, Germany

**Author notes:** Corresponding author: Wim Versées, **Email:**.

**Keywords:** Parkinson’s disease, LRRK2, drug design, allosteric inhibitor, Nanobody

## Abstract

Mutations in the gene coding for Leucine-Rich Repeat Kinase 2 (LRRK2) are a leading cause of the inherited form of Parkinson’s disease (PD), while LRRK2 overactivation is also associated with the more common idiopathic form of PD. LRRK2 is a large multi-domain protein, including a GTPase as well as a Ser/Thr protein kinase domain. Common disease-causing mutations increase LRRK2 kinase activity, presenting LRRK2 as an attractive target for inhibitory drug design. Currently, drug development has mainly focused on ATP-competitive kinase inhibitors. Here, we report the identification and characterization of a variety of Nanobodies that bind to different LRRK2 domains and inhibit or activate LRRK2 activity in cells and *in vitro.* Importantly, diverse groups of Nanobodies were identified that inhibit LRRK2 kinase activity through a mechanism that does not involve binding to the ATP pocket or even to the kinase domain. Moreover, while certain Nanobodies completely inhibit the LRRK2 kinase activity, we also identified Nanobodies that specifically inhibit the phosphorylation of Rab protein substrates. Finally, in contrast to current type-I kinase inhibitors, the studied kinase-inhibitory Nanobodies did not induce LRRK2 microtubule association. These comprehensively characterized Nanobodies represent versatile tools to study the LRRK2 function and mechanism, and can pave the way toward novel diagnostic and therapeutic strategies for PD.

## Introduction

Parkinson’s disease (PD) is a common and devastating neurodegenerative movement disorder affecting around 2 % of the global population (1). The disease is characterized by degeneration of dopaminergic neurons, which leads to the typical symptoms including resting tremor, bradykinesia and postural instability. Although treatments to alleviate these symptoms have been available since long, there is still no cure. A very promising strategy that is currently being intensively pursued is the targeting of the protein Leucine-Rich Repeat Kinase 2 (LRRK2). Mutations in LRRK2 are among the most common causes of familial PD (2), while an increased LRRK2 activity has also been associated with the more frequent idiopathic form of PD (3, 4). Moreover, LRRK2 mutations and/or overexpression have also been linked to a number of chronic inflammatory conditions, including Crohn’s disease (5, 6).

LRRK2 is a large multi-domain protein belonging to the ROCO family (Fig. 1a), that bears a rather unique combination of two catalytic activities: GTPase activity mediated by the Roc domain and Ser/Thr protein kinase activity (7). Recently, several Rab GTPases were identified as the physiological substrates of LRRK2 kinase activity (8, 9), while LRRK2 is also known to auto-phosphorylate (10). Although the details of the regulatory mechanism of LRRK2 are not yet completely understood, we previously showed, using a more simple LRRK2 homologue from the bacterium *Chlorobium tepidum* (CtRoco), that the RocCOR supra-domain undergoes a dimer-monomer cycle concomitant with GTP binding and hydrolysis (11). This is in line with findings for LRRK2 in cells, which show that the protein predominantly occurs as a monomer with low kinase activity in the cytosol and as a dimer with high kinase activity at the membrane (12, 13). These results point toward a complex interplay between the GTPase and kinase domains of LRRK2, regulated by large-scale conformational changes.

**Figure 1:**
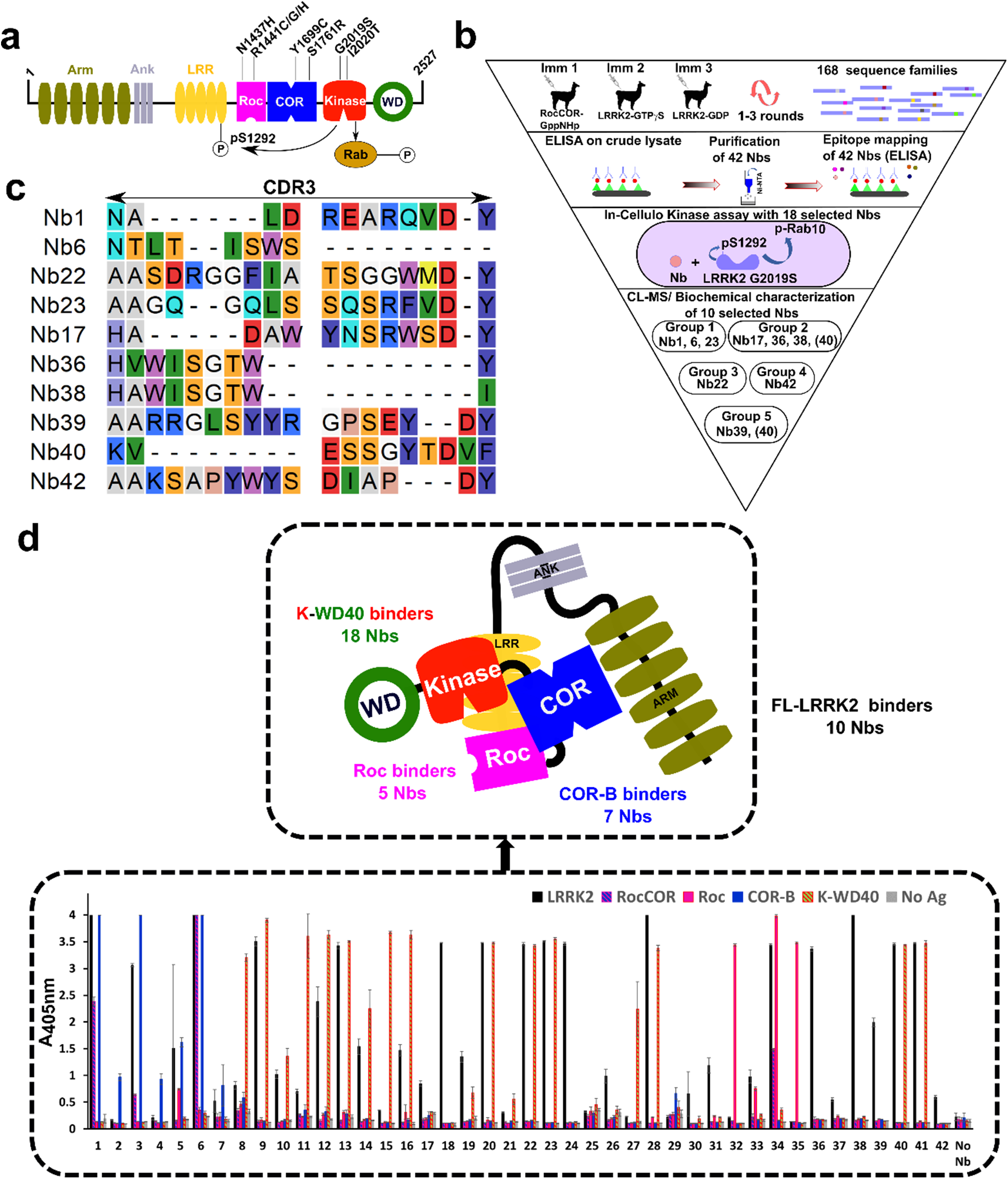
Identification of LRRK2-targeting Nanobodies (Nbs). **(a)** Domain arrangement of LRRK2, with important PD mutations indicated. Two LRRK2 kinase activities relevant to this study are also indicated: phosphorylation of Rab proteins, and autophosphorylation at position S1292. **(b)** Funnel approach used in this study to identify and characterize LRRK2-binding and modulating Nbs. **(c)** Sequences of the CDR3 regions of the 10 Nbs that were analyzed in detail. Nb36 and Nb38 belong to the same CDR3 sequence family. **(d)** Domain mapping of the purified Nbs using ELISA on either full-length LRRK2 (FL-LRRK2) or the RocCOR, Roc, C-terminal part of COR (COR-B) or kinase-WD40 (K-WD40) constructs (*lower panel*). Each ELISA signal is the average of three experiments. This domain mapping reveals 7 Nbs that bind on COR-B, 5 Nbs that bind on Roc, 18 Nbs that bind on K-WD40, and 10 Nbs that only bind on FL-LRRK2 (*upper panel*).

The most prevalent PD mutations in LRRK2 are clustered within the catalytic RocCOR and kinase domains (Fig. 1a), and several PD mutations lead to a decrease in GTPase and/or an increase in kinase activity (14, 15). Most notably, autophosphorylation of Serine-1292 (16) and Rab protein phosphorylation (8) are increased by pathogenic LRRK2 variants, and particularly by the most common G2019S mutation. These findings support the idea that LRRK2 mutations cause PD through a gain-of-function mechanism, and inhibition of LRRK2 kinase activity is therefore considered to be a particularly promising strategy for the treatment of PD (17, 18). However, preclinical studies have indicated that long-term inhibition of LRRK2 with ATP-competitive inhibitors might potentially be associated with toxic side effects, including kidney abnormalities in rodents and an accumulation of lamellar bodies in type II pneumocytes in the non-human primate lung (19–21). Targeting the multiple enzymatic functions and regulatory mechanisms of LRRK2 in an allosteric way, using compounds that bind outside the ATP pocket, could form a very attractive alternative to currently explored approaches (22). One way to modulate the dynamics, regulation and activity of proteins is via the use of Nanobodies (Nbs), the small and stable single-domain fragments derived from camelid heavy chain–only antibodies (23). Accordingly, we recently reported the identification of Nbs that allosterically activate the GTPase activity of a bacterial LRRK2 homologue by interfering with the protein’s monomer-dimer cycle (24).

Here, we report the identification and *in vitro* and *in cellulo* characterization of Nbs acting as allosteric modulators of human LRRK2 kinase activity. A wide range of Nb families are discovered that bind to different LRRK2 domains with a variety of affinities. Several of these Nbs robustly inhibit LRRK2 kinase activity, both in cells and *in vitro*, while others significantly activate LRRK2 activity. Moreover, within the set of LRRK2-inhibiting Nbs, different modes-of-action can be discerned with some inhibiting total kinase activity, while others specifically inhibit phosphorylation of the Rab substrates. Interestingly, a subset of the Nbs inhibit kinase activity while not binding directly on the kinase domain and act as mixed (non-competitive)-type inhibitors, showing that the Nbs use an allosteric mechanism of inhibition. Finally, we show that in contrast to currently available ATP-competitive kinase inhibitors, the Nbs do not induce formation of LRRK2 filaments on microtubules in cells. This study thus proposes these Nbs as novel inhibitors of LRRK2 kinase activity that act via a completely different mode-of-action compared to the currently available ATP-competitive kinase inhibitors. Thus, this finding may open new therapeutic strategies in the fight against Parkinson’s disease.

## Results

### Identification of a wide variety of LRRK2-binding Nbs

To identify Nbs that bind LRRK2 and to maximize chances of finding LRRK2 activity-modulating Nbs, three immunization strategies were followed, each time using a different llama (Fig. 1b). In a first immunization strategy we immunized a llama with the LRRK2 RocCOR construct (Fig. S1), and after immunization Nbs were selected via a phage display panning approach using full-length LRRK2 as the bait protein. In immunization strategies 2 and 3, we immunized llamas with LRRK2 either in presence of a large excess of GTPγS (a non-hydrolysable GTP analogue) or in presence of a large excess of GDP, and phage display selections were performed using the corresponding proteins. Additionally, to enrich for Nbs binding to the Roc domain, phage display selections were performed using either the GTPγS- or GDP-bound Roc domain protein. These different strategies finally resulted in 49 Nb families originating from immunization 1, 70 Nb families originating from immunization 2 with selection against LRRK2-GTPγS, 4 Nb families originating from immunization 2 with selection against Roc-GTPγS, 44 Nb families originating from immunization 3 with selection against LRRK2-GDP, and 1 Nb family originating from immunization 3 with selection against Roc-GDP (each Nb family displaying a unique complimentary determining region 3 (CDR3) sequence). Based on an initial ELISA screening on crude *E. coli* cell extracts expressing these Nbs, 42 Nbs belonging to 41 different CDR3 sequence families were selected, expressed and purified to homogeneity (Fig. 1c, Table S1, Fig. S2).

Subsequently, binding to full-length LRRK2 was confirmed for this set of purified Nbs using ELISA, while in parallel also a first mapping of the binding regions of these Nbs was performed by assessing binding to purified RocCOR, Roc, COR-B and kinase-WD40 (K-WD40) domains of LRRK2 (Fig. 1d). Nearly all tested Nbs show binding to full-length LRRK2 and/or at least one of the domain constructs, except for two Nbs (Nb25 and Nb29). All 7 purified Nbs that resulted from the immunization with the RocCOR domain construct (Immunization 1) specifically interacted with the C-terminal subdomain of the COR domain (COR-B). Among the Nbs obtained from the immunizations and selections using full-length LRRK2, the majority bound within the K-WD40 domain of the protein (18 Nbs). Another subset of 10 Nbs showed robust binding to LRRK2 while no binding was observed to any of the individual LRRK2 domain constructs. We therefore assume that these Nbs either bind exclusively to the N-terminal region of LRRK2 (armadillo-ankyrin-LRR domains) that was not covered by individual domains in the ELISA, and/or bind on an epitope on the interface of two or more domains and thus require the full-length LRRK2 protein for binding. Finally, a last set of Nbs were specifically selected by phage display panning on the LRRK2 Roc domain, resulting in 5 Roc domain-binding Nbs (Nb32, 33, 34, 35, 42; note that Nb42 did not show clear binding to Roc in ELISA but binding was confirmed by size exclusion chromatography, see Fig. S3). The results of this initial domain mapping are schematically summarized in Fig. 1d.

### Nbs bind LRRK2 and modulate its activity in cells

After confirming binding of 40 purified Nbs to LRRK2 *in vitro*, we next wanted to confirm whether these Nbs bound to LRRK2 in human HEK293 cells over-expressing LRRK2. Hereto, we selected a subset of 18 Nbs, considering Nbs originating from the three immunization strategies and targeting different LRRK2 domains. The Nbs were expressed as GFP-fusions from a pEGFP vector in LRRK2(wild-type)-overexpressing HEK293 cells, and a pull-down experiment was performed using magnetic GFP-nanotrap beads (Fig. 2a). This experiment showed that all tested Nbs were able to pull-down LRRK2 under these conditions. This thus indicates that these 18 Nbs are functional as intrabodies in the context of the cytoplasm of human cells and have sufficiently high affinity to pull-down their target protein.

**Figure 2:**
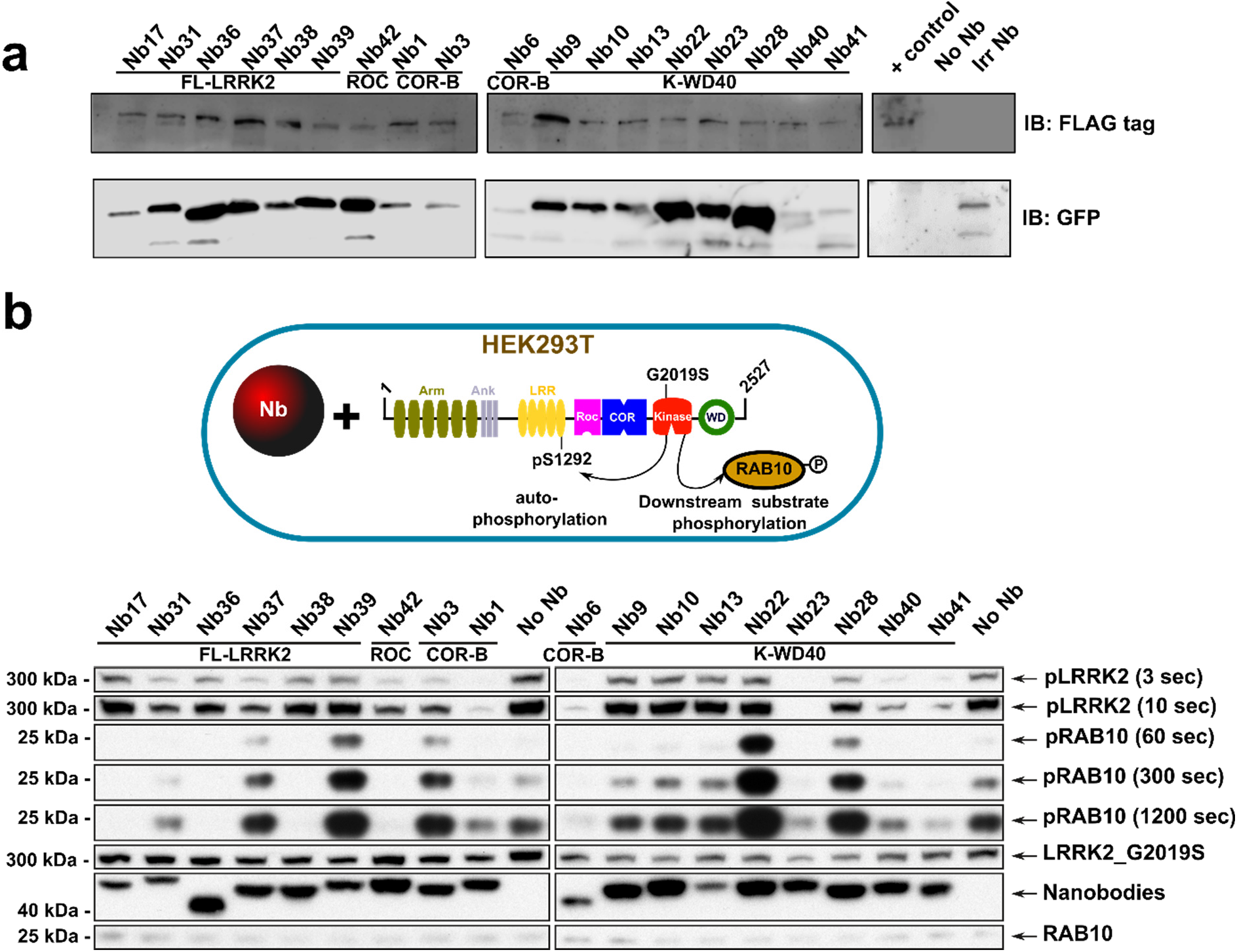
Nbs engage LRRK2 and modulate LRRK2 activity in cells. **(a)** Nbs bind and immuno-precipitate LRRK2. Pull-down assays after transient co-transfection of HEK293 cells with GFP-tagged Nbs and (S)trep-(F)lag-tagged LRRK2, using magnetic GFP-Trap beads. HEK293 cell lysate only overexpressing SF-LRRK2 (“No Nb”) or expressing an irrelevant Nb (“Irr Nb”) were used as negative controls. LRRK2 (*upper row*) and GFP-Nbs (*lower row*) were detected via immunoblotting. Blot is representative of n=3. **(b)** Influence of Nbs on the kinase activity of the LRRK2(G2019S) variant in HEK293T cells. LRRK2(G2019S) and its effector Rab29 were overexpressed together with GFP-tagged Nbs in HEK293T cells. A negative control, where no Nb is overexpressed (“No Nb”) is also included. In rows labeled “pLRRK2”, LRRK2 pS1292 levels are determined by Western Blot using a site-specific anti-pLRRK2(pS1292) antibody (Abcam, ab203181) (shown at different times of development). In the rows labeled “pRAB10”, endogenous pT73-Rab10 levels are determined by Western Blot using the MJFF/Abcam antibody MJF-R21 (Abcam, ab230261) (shown at different times of development). The three lower rows contain controls of LRRK2 (rat anti-LRRK2, 24D8), GFP-Nb (rat anti-GFP) and Rab10 (rabbit anti-Rab10) expression levels, determined on a different Western Blot than pLRRK2 and pRab10. Blot is representative of n=3 (see Fig. S4).

Subsequently, to assess the influence of these 18 Nbs on in-cell LRRK2 kinase activity, we monitored two activities of LRRK2: phosphorylation of the endogenous substrate Rab10 at position T73 and LRRK2 autophosphorylation at position S1292 (8, 9, 16, 25). Both activities have previously been shown to be increased in the most relevant LRRK2 PD-mutants, including the common G2019S mutant. In this case, we thus co-expressed the LRRK2(G2019S) mutant together with the different Nb-GFP fusions in HEK293T cells. Among the 18 selected Nbs, seven Nbs (Nb3, Nb9, Nb10, Nb13, Nb31, Nb37, Nb39) did not consistently and/or considerably influence LRRK2 kinase activity in comparison to the negative control (Fig. 2b; Fig. S4). Four Nbs strongly decreased both LRRK2 autophosphorylation and Rab10 phosphorylation in cells: Nb1, Nb6, Nb23 and Nb42. Interestingly, another five inhibitory Nbs only decrease the level of Rab10 phosphorylation, while they largely leave autophosphorylation unaffected: Nb17, Nb36, Nb38, Nb40 and Nb41. Finally, two Nbs increase the levels of LRRK2-mediated Rab10 phosphorylation: Nb22 and Nb28. Based on these results and their inhibitory/activating effect on LRRK2 kinase activity, we selected the following ten Nbs for further in-depth characterization: Nb1, Nb6, Nb17, Nb22, Nb23, Nb36, Nb38, Nb39, Nb40, Nb42 (Fig. 1c).

### Epitope mapping and affinity of LRRK2 modulating Nbs

In addition to the ELISA experiment described above (Fig 1d), we used a crosslinking mass spectrometry (CL-MS) approach to obtain more detailed insight in the binding epitopes of the ten selected Nbs. The CL-MS data revealed that the different Nbs crosslink to lysines within LRRK2 predominantly via a conserved lysine residue in the framework 3 region of the Nb. This lysine residue is present in all selected Nbs except for Nb6, for which correspondingly no crosslinking data could be obtained. Overall, the CL-MS data are in very good agreement with the results of the domain mapping using ELISA (Fig. 3a). Based on the ELISA experiments, Nb17, Nb36, Nb38 and Nb39 were found to bind either on the N-terminal domains of LRRK2 or to bind on the interface of several domains (“full-length LRRK2 binders”). Correspondingly, CL-MS reveals that all these Nbs make multiple contacts to the N-terminal LRR domain as well as C-terminal parts of LRRK2, including the C-terminal end of COR (Nb39), the kinase domain (Nb17 and Nb38) and both the kinase and WD40 domain (Nb36). For Nb42 only one crosslink with a lysine (K1502) within the Roc domain is identified, in good agreement with the domain mapping in ELISA. Nb22, Nb23 and Nb40 were all identified as kinase-WD40 binders in ELISA, and correspondingly the crosslinking data reveals interactions with the WD40 domain (Nb22 and Nb40) or the kinase domain (Nb23). Interestingly, Nb1 which was unequivocally identified as a COR-B binder in ELISA, correspondingly makes multiple crosslinks with the COR-B domain (K1824, K1832, K1833), but also crosslinks are found with residues within other LRRK2 domains, including the kinase domain and the leucine-rich repeats. This finding is in good agreement with a very central localization of the COR-B domain within recent structural models of LRRK2 (26–29).

**Figure 3:**
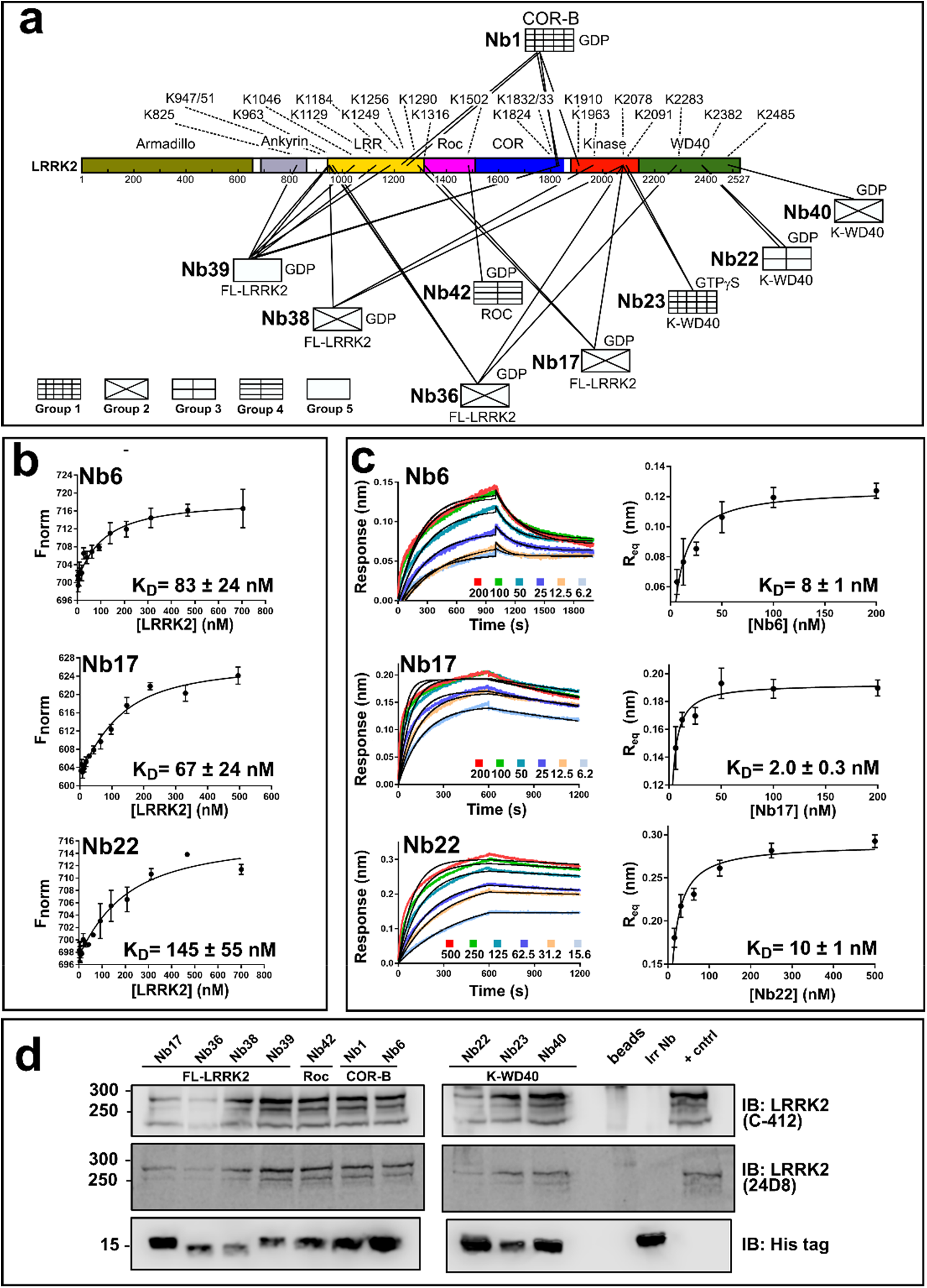
Nbs bind LRRK2 with high affinity via interactions with different domains. **(a)** Mapping of the Nb binding epitopes using CL-MS. The Nbs are divided into 5 groups according to their effect on LRRK2 kinase activity, as defined in Fig. 1b. The observed cross-links between the Nbs and LRRK2 are indicated by lines, with the corresponding lysine residues on LRRK2 indicated by their residue number. The domain specificity of the Nbs as determined in ELISA is given below the respective Nbs. **(b)** Affinity measurements of the Nbs for LRRK2 using MST. Binding isotherms shown for three representative Nbs, Nb6 (group1), Nb17 (group2) and Nb22 (group3) (see Fig. S5 for all curves), were obtained by titrating increasing concentrations of LRRK2 to fluorescently labeled Nbs. The K_D_ values (± standard error) from fitting on a quadratic binding equation are given (each data point is the average ± SD of n=3). **(c)** Affinity measurements of the Nbs for LRRK2 using BLI. BLI sensorgrams for a dilution series of the Nb, with Nb concentrations as indicated on the figure (*left*), and the resulting dose-response curves (*right*) are shown for the same 3 Nbs as in (b) (see Fig. S6 for all curves). The K_D_ values (± standard error) obtained by fitting the dose-response curves with a Langmuir binding equation are given (each data point is the average ± SD of n=3). **(d)** Nbs immuno-precipitate endogenous (mouse) LRRK2. Lysates derived from RAW264.7 cells were incubated with 1.5µM of purified His-tagged Nbs and pull-downs were performed using magnetic Dynabeads. LRRK2 was detected via immunoblotting, using two different antibodies (C-412 and 24D8). The blot is representative of n=3.

To determine the binding affinity (dissociation constant K_D_) of the ten Nbs, two methods were used in parallel: microscale thermophoresis (MST) and bio-layer interferometry (BLI). For the MST experiment, we site-specifically labeled the ten Nbs with an m-TAMRA fluorophore at its C-terminus, using sortase-mediated coupling (24), and then titrated increasing amounts of full-length LRRK2 to these Nbs (Fig. 3b, Fig.S5). For all 10 Nbs an MST signal was observed, except for Nb23, which did not generate a change in thermophoresis behavior upon binding to LRRK2. For the other Nbs, K_D_ values are found in the range of 25nM-150nM (Table 1). For the BLI experiment LRRK2 was first trapped on a Streptavidin-coated biosensor using biotinylated Nb40 as trapping agent (except to assess binding of Nb40, where Nb42 was used as trapping agent), after which binding of all Nbs to LRRK2 was determined. A clear binding signal was obtained for all Nbs (including Nb23), with some Nbs showing an indication of a second slow phase in the binding and dissociation curves, which could either reflect conformational changes upon Nb binding or the presence of pre-existing conformational heterogeneity in LRRK2. Fitting of the sensorgrams was performed initially using a 1:1 binding model, and the resulting R_eq_ values were subsequently used to calculate K_D_ values from the corresponding dose-response curves fitted on a Langmuir model. This yields K_D_ values ranging from 2 nM – 60 nM (Fig. 3c; Fig. S6). Overall, a (nearly 5 to 10-fold) higher affinity is obtained with BLI compared to MST (Table 1). These differences are probably due to the experimental set-up, with MST using LRRK2 in solution and BLI using LRRK2 trapped on a surface by means of a second Nb, which might also result in different LRRK2 conformations. Additionally, the K_D_ values obtained via the MST titration will scale with the concentration of properly folded LRRK2 in the sample during the measurement, and denaturation of a part of the LRRK2 protein could hence lead to an overestimation of the K_D_ values (note that in MST LRRK2 is used at varying concentrations, while in BLI the Nbs are used at varying concentrations). Subsequently, a BLI experiment was performed using the biotinylated Roc-COR-kinase-WD40 (RCKW) construct of LRRK2 coupled to the Streptavidine-coated biosensors and titration with increasing amount of Nbs (Fig. S7). In agreement with the ELISA and CL-MS experiments, which showed that Nb17, Nb36, Nb38 and Nb39 interact with LRRK2 N-terminal domains and thus require full-length LRRK2 for binding, no binding to the RCKW construct is observed for these Nbs. Comparison of the K_D_ values obtained with BLI of Nb1, Nb6, Nb22 and Nb40 for full-length LRRK2 and the RCKW protein (Table 1), shows that these values are in relatively good agreement, indicating that the N-terminal domains of LRRK2 only play a minor role in the equilibrium binding affinities of those Nbs.

**Table 1:** Equilibrium dissociation constants (K_D_ in nM) for binding of the set of 10 Nbs (belonging to 5 functional groups based on their effect on kinase activity) to either full-length LRRK2 or its RCKW domain construct, as assessed by two methods in parallel: microscale thermophoresis (MST) and biolayer interferometry (BLI).

| Nanobody | Binding epitope (ELISA) | Nb group | Full-length LRRK2 $K_D$ (nM) | | RCKW $K_D$ (nM) |
| --- | --- | --- | --- | --- | --- |
|  |  |  | MST | BLI | BLI |
| **Nb1** | COR-B | Group 1 | $91 \pm 28$ | $55 \pm 12$ | $39 \pm 4$ |
| **Nb6** | COR-B | | $83 \pm 24$ | $8 \pm 1$ | $5 \pm 1$ |
| **Nb23** | K-WD40 | | NBS<sup>a</sup> | $20 \pm 5$ | $7 \pm 4$ |
| **Nb17** | FL-LRRK2<sup>c</sup> | Group 2 | $67 \pm 24$ | $2.0 \pm 0.3$ | NB<sup>b</sup> |
| **Nb36** | FL-LRRK2<sup>c</sup> | | $78 \pm 25$ | $16 \pm 8$ | NB<sup>b</sup> |
| **Nb38** | FL-LRRK2<sup>c</sup> | | $48 \pm 11$ | $37 \pm 4$ | NB<sup>b</sup> |
| **Nb40** | K-WD40 | | $26 \pm 10$ | $2.5 \pm 0.3$ | $2.6 \pm 0.3$ |
| **Nb22** | K-WD40 | Group 3 | $145 \pm 55$ | $10 \pm 1$ | $5 \pm 1$ |
| **Nb42** | Roc | Group 4 | $94 \pm 30^d$ | $58 \pm 18$ | $18 \pm 2$ |
| **Nb39** | FL-LRRK2<sup>c</sup> | Group 5 | $79 \pm 19$ | $12 \pm 3$ | NB<sup>b</sup> |
| <sup>a</sup> NBS: no binding signal observed in MST<br><sup>b</sup> NB: no binding on RCKW<br><sup>c</sup> FL-LRRK2: full-length LRRK2<br><sup>d</sup> Value determined in presence of GTPγS instead of GDP, which was used in all the other measurements | | | | | |

Considering their high affinity binding, we next tested whether those Nbs would also be able to pull-down LRRK2 at endogenous/physiological expression levels. Therefore, we turned to lysates of mouse RAW264.7 cells, which express LRRK2 at relatively high levels (30). Interestingly, we find that all 10 Nbs efficiently pull-down LRRK2 when added to these lysates (Fig. 3d), showing that (1) the Nbs are cross-reactive toward mouse LRRK2, and (2) their affinity is sufficiently high to pull-down endogenous levels of LRRK2 from cell lysates.

### Nbs inhibit LRRK2 peptide and/or Rab phosphorylation *in vitro*

To test if the 10 Nbs directly influence LRRK2 (wild-type) kinase activity, we next performed *in-vitro* assays using purified proteins. In a first approach we measured LRRK2 kinase activity vis-à-vis an optimized LRRK2 peptide substrate (AQT0615) using the commercially available PhosphoSens® Protein Kinase Assay (AssayQuant Technologies Inc.) (Fig. 4a), while in a second approach we also measured the influence of the Nbs on LRRK2-mediated Rab8a phosphorylation using a Western blot-based method (Fig. 4b). The influence of all 10 Nbs on LRRK2 activity was screened at a fixed Nb concentration of 25 µM, and compared to a no-Nb negative control, and a positive control where the ATP-competitive LRRK2 inhibitor MLi-2 was added (31–33). These experiments were performed with LRRK2 either in the presence of a large excess (500µM) of GDP or GTPγS, but no significant influence of the nucleotide on the inhibition profile was observed (Fig. S8). Consistent with the *in cellulo* data, the WD40-binding Nb22 activates the kinase activity of LRRK2, both toward the peptide and Rab8a substrate, by about 30 to 50 % compared to the controls. The group of Nbs which showed an influence on Rab phosphorylation in cells while leaving autophosphorylation unaffected (Nb17, Nb36, Nb38 and Nb40; Fig. 2b), very consistently only show a very small or no effect on LRRK2 peptide phosphorylation, while they have a prominent effect on Rab8a phosphorylation *in vitro* (although less pronounced for Nb40).

**Figure 4:**
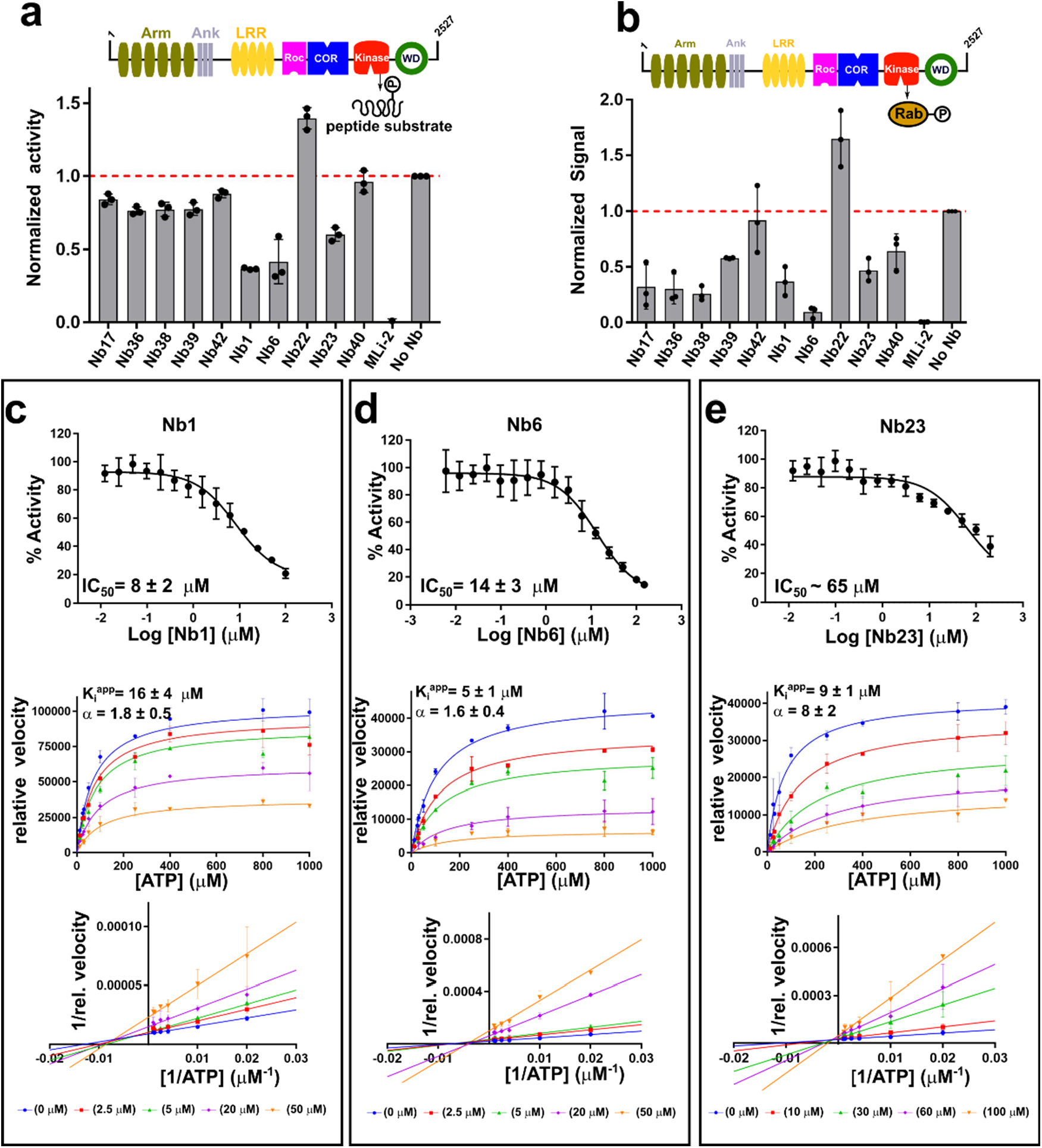
Modulation of *in vitro* kinase activity by LRRK2-targeting Nbs. **(a)** Effect of Nbs on LRRK2 kinase activity measured using the LRRK2-optimized AQT0615 peptide as substrate. (**b**) Effect of Nbs on LRRK2-mediated phosphorylation of Rab8a determined via a Western blot assay (see Fig. S8). In both (a) and (b) the influence of the Nbs (25 µM) on the relative kinase activity compared to the “No-Nb” control is plotted, and a positive control with 0.2 µM MLi-2 is included. Each bar reflects the average (± SD) of three independent measurements. (**c-e**) Dose-response curves (*upper panels*) for the inhibition of the *in vitro* LRRK2 kinase activity by the group1 Nbs: Nb1 (**c**), Nb6 (**d**) and Nb23 (**e**), using a serial dilution of the Nb and a fixed concentration of AQT0615 peptide (10µM) and ATP (1 mM). The *middle panels* show the Michaelis-Menten curves obtained for LRRK2 at varying concentrations of ATP and a fixed (sub-saturating) concentration of peptide substrate (AQT0615) and at varying concentrations of the respective Nbs. The *lower panels* show the corresponding linearizations according to the Lineweaver-Burk method (double-reciprocal plot). The Nb concentrations used are indicated below the plots. Each datapoint reflects the average (± standard error) of three independent measurements. The IC_50_ (± SD) values resulting from fitting on a three-parameter logistic equation and the K_i_^app^ and α values (± SD) resulting from global fitting on a mixed-type inhibition mechanism are indicated on the graphs.

Combined, our *in cellulo* and *in vitro* assays, thus suggest that these four Nbs do not directly affect LRRK2 kinase activity *per se*, but rather specifically influence Rab phosphorylation. In contrast, Nb1, Nb6 and (to a lesser extent) Nb23 have a clear inhibitory effect on LRRK2 kinase activity toward both peptide and Rab substrates *in vitro*. This is again in very good agreement with our *in cellulo* data (Fig. 2b) which showed that these Nbs inhibit both LRRK2 autophosphorylation and Rab phosphorylation, thus suggesting that these Nbs act as true inhibitors of total LRRK2 kinase activity. One exception seems to be Nb42, which inhibited both LRRK2 autophosphorylation and Rab phosphorylation in cells, while no inhibitory effect could be observed *in vitro*.

Since we found that Nb1, Nb6 and Nb23 inhibit LRRK2 kinase activity with respect to peptides and Rab proteins *in vitro*, we continued to perform a dose-response analysis with these three Nbs using the same model peptide substrate and at a fixed concentration of 1 mM of ATP (Fig. 4c-e). Fitting of these dose-response curves yielded IC_50_ values of 8 ± 2 µM and 14 ± 3 µM for Nb1 and Nb6, respectively, and an approximate IC_50_ value of 65 µM for Nb23. Similarly, a dose response was observed for Nb17, Nb36 and Nb38 with respect to LRRK2 Rab8a phosphorylation (Fig. S8), but saturation of the blot at high Nb concentrations prevented a more qualitative treatment of these curves.

### LRRK2-inhibiting Nbs act via a non-ATP competitive / allosteric mechanism

Three Nbs (Nb1, Nb6, Nb23) were identified that robustly inhibit LRRK2 autophosphorylation and Rab phosphorylation activity in cells, as well as LRRK2 kinase activity towards peptide and Rab substrates *in vitro*. Combined these experiments suggest that these Nbs target the LRRK2 kinase activity *per se*. Interestingly, while the ELISA and CL-MS experiments suggest that Nb23 binds to the kinase domain, Nb1 and Nb6 bind on the COR-B domain. This strongly suggests that at least the latter two Nbs act as allosteric kinase inhibitors. To further confirm this, we set out to determine the mechanism of inhibition (competitive vs. uncompetitive vs. mixed/non-competitive) of these three Nbs with regard to ATP. Full Michaelis-Menten curves were obtained at a fixed (probably sub-saturating) concentration of peptide substrate (10 µM) and varying concentrations of ATP (Fig. 4c-e). For Nb1 and Nb6, linearization of the curves using the Lineweaver-Burk plot clearly shows intersecting lines left of the Y-axis, indicative of a mixed-type inhibition. This confirms that these Nbs are not competing with ATP for binding and that they inhibit the reaction by binding on an allosteric site, in agreement with the ELISA and CL-MS domain/epitope mapping data. For Nb1, a global fit of the kinetic data using a mixed inhibition model accordingly gives a K_i_^app^ of 16 ± 4 µM with an α-value of 1.8 ± 0.5, corresponding to a K_ic_^app^ (= apparent K_i_ for apo-LRRK2) of 16 µM and a K_iu_^app^ (= apparent K_i_ for ATP-bound LRRK2) of 30 µM. For Nb6, fitting on the same model yields a K_i_^app^ of 5 ± 1 µM with an α-value of 1.6 ± 0.4, corresponding a K_i_^app^ of 5 µM and a K_iu_^app^ of 8 µM. For the kinase domain-binding Nb23, the linearized curves intersect closer to the Y-axis, indicating a mechanism which is more ATP-competitive like. Yet, the lines do not exactly intersect on the Y-axis and a systematic decrease in V_max_^app^ and increase (or no effect) in K_M_^app^ value with increasing Nb concentration is observed, which also here indicates a mixed-inhibition mechanism (Fig. S9). Fitting on a mixed inhibition model for Nb23 gives a K_i_^app^ of 9 ± 1 µM with an α-value of 8 ± 2, corresponding a K_ic_^app^ of 9 µM and a K_iu_^app^ of 72 µM.

To further confirm that Nb1, Nb6 and Nb23 bind outside the kinase ATP-binding pocket and thus target a different site than the “classical” ATP-competitive inhibitors, such as MLi-2 (31–33), we performed a competition ELISA titration experiment (Fig. S10). In this experimental setup, the ELISA experiment was performed using a fixed concentration of LRRK2 coated on the bottom of the ELISA plate, and using a dilution series of either of the three Nbs, resulting in a dose-response titration curve reflecting an apparent affinity of the Nbs. Subsequently, a repetition of the setup in the presence of a large excess of MLi-2 (1 µM), or of the corresponding non-tagged Nb (at 9 µM) as a positive control, was performed. While a very prominent rightward shift in the titration curves is observed when adding the corresponding untagged Nb as a direct orthosteric competitor, no rightward shift is observed for any of the three Nbs in presence of MLi-2. This thus further proves that neither of these three Nbs bind in the same pocket as the Type I ATP-competitive inhibitor MLi-2 (31–34), thus confirming that these Nbs act via a non-ATP competitive allosteric inhibitory mechanism.

Together, our findings that (i) Nb1, Nb1 and Nb23 act as mixed-type inhibitors of LRRK2 kinase activity with respect to ATP, and that (ii) the binding of these Nbs is not outcompeted by a large excess of the ATP pocket-binding inhibitor Mli-2 (31–34), shows that these Nbs act as non-ATP competitive inhibitors and target a site different from the ATP pocket. Nb1 and Nb6 moreover bind to an entirely different domain (COR-B), further illustrating their allosteric mode of inhibition (i.e. allosteric with regard to the ATP binding pocket).

### Expression of LRRK2-targeting Nbs did not result in LRRK2 relocalization to microtubules

LRRK2 pharmacological kinase inhibitors of different structural classes induce cellular recruitment of overexpressed LRRK2 to microtubules, similar to four out of the five major PD-causing mutations (35, 36). The binding of LRRK2 to microtubules subsequently reduces the kinesin- and dynein-mediated transport along microtubules (27). In order to investigate whether our ten identified LRRK2 kinase-modulating Nbs result in a similar phenotype, HEK293 cells were co-transfected with constructs coding for mScarlet-LRRK2 and GFP-Nbs. For all 10 Nbs analyzed, confocal microscopy analysis shows that no relocalization to microtubules is observed 48h after co-transfection with the Nbs, in contrast to MLi-2 treatment where filamentous skein-like structures of LRRK2 are observed, indicating that the Nbs trap LRRK2 in a different conformation compared to classical inhibitors (Fig. S11). Next, we wanted to investigate whether the LRRK2-targeting Nbs have the ability to alter the recruitment of LRRK2 to microtubules induced by inhibition of LRRK2 kinase activity by the specific ATP-competitive inhibitor MLi-2 (31–34). To do that, HEK293 cells co-transfected with mScarlet-LRRK2 and GFP-Nb constructs were treated with 1 µM of MLi-2 (Fig.5, Fig. S12). Interestingly, cells co-transfected with a subset of the tested Nbs showed no MLi-2-induced recruitment to microtubules, and LRRK2 cytoplasmic distribution was maintained in contrast to controls with MLi-2-treated cells co-transfected with GFP or an irrelevant Nb. This rescuing effect was most prominent for Nb17, Nb22, Nb23 and Nb40. While Nb36 and Nb38 also significantly reduced the number of cells with skein-like structures, some shorter filaments are still present in some cells transfected with these two Nbs.

**Figure 5:**
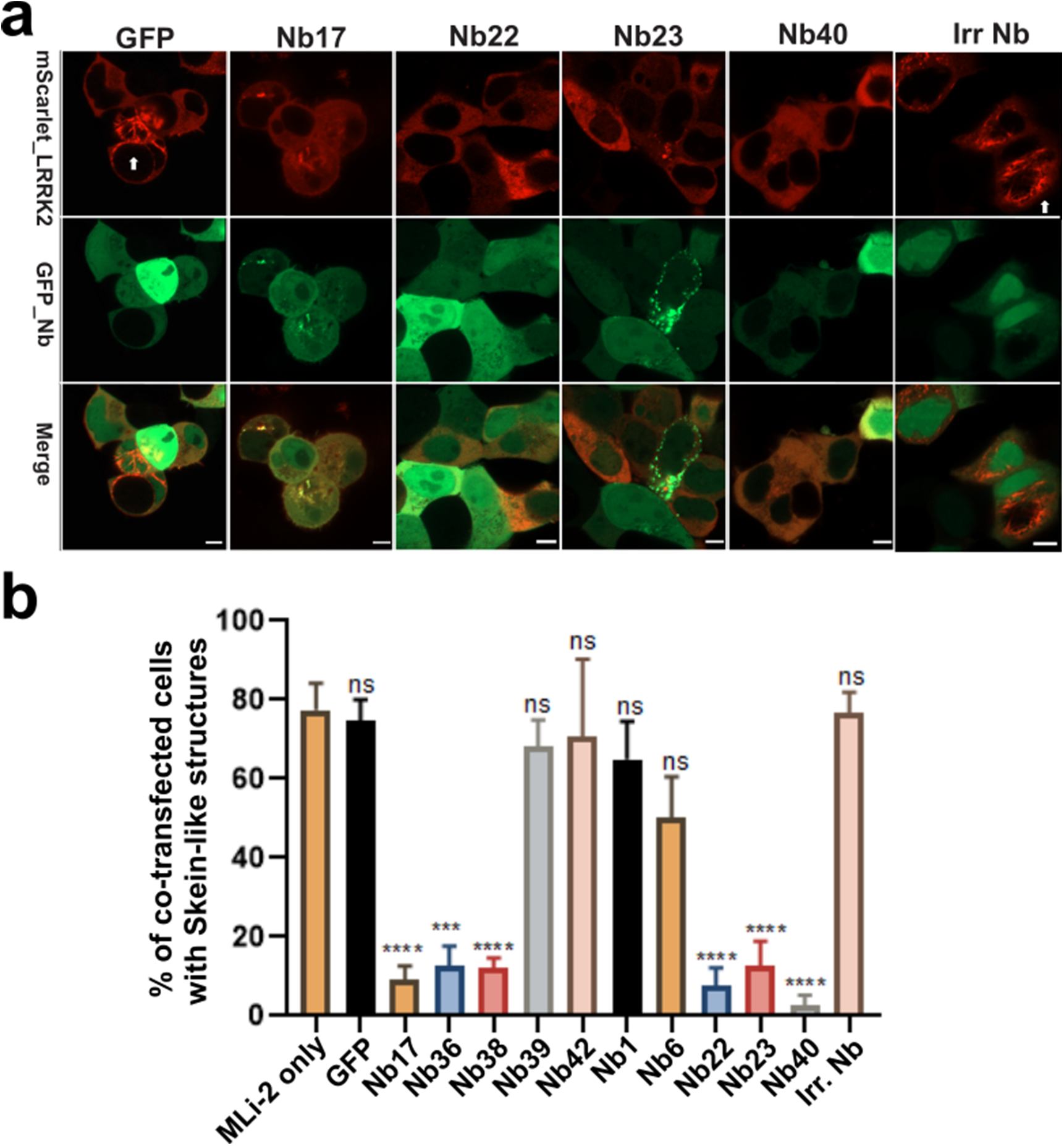
Nbs can rescue MLi-2 induced LRRK2 relocalization to microtubules. **(a)** Effect of Nbs on MLi-2-induced microtubule relocalization of LRRK2 (results for a selected set of Nbs are shown, see Fig. S12 for all data). HEK293 cells were co-transfected with GFP or the GFP-tagged Nbs and mScarlet-LRRK2 for 24hrs and then treated with 1 µM MLi-2 for 90mins. Co-transfection of mScarlet-LRRK2 with only GFP or an irrelevant control Nb show the MLi-2-induced filamentous skein-like structures of LRRK2 indicated by white arrows. Co-transfection with a subset of GFP-Nbs inhibited the MLi-2-induced LRRK2 relocalization. Scale bar = 5 µM. (**b**) Quantification of results shown in (a) and Fig.S12. A minimum of 200 transfected cells were analyzed under each condition for filamentous structures of GFP-LRRK2. The average percentage of cells showing typical skein-like structures and standard errors of the mean (SEM) for three biological replicates are shown with p values: One-way ANOVA and Dunnett’s multiple comparisons test (MLi-2 only as a control), **** p ≤ 0.0001, *** p ≤ 0.001, ns: not significantly different (p > 0.05).

## Discussion

LRRK2 is an intensively pursued target for drug development in the fight against PD, and might also hold promise to treat certain inflammatory diseases such as Crohn’s disease (6, 17, 18, 31). However, certain concerns for potential adverse side effects associated with the use of small-molecule kinase inhibitors directed toward the LRRK2 ATP-binding pocket remain (19, 20, 37). Here, we report the identification of a large and diverse repertoire of Nbs directed toward different domains of LRRK2 (Fig. 1). After a pre-selection based on their effect on LRRK2 kinase activity in cells, we chose to characterize ten of these Nbs in detail. All ten Nbs were able to efficiently pull-down LRRK2 from cell lysates, both upon LRRK2 over-expression and at endogenous LRRK2 expression levels. This reflects a high affinity binding, which was subsequently confirmed using a combination of two biophysical methods which yielded K_D_ values ranging from 2 to 150 nM.

We analyzed the influence of this set of Nbs on LRRK2 autophosphorylation and Rab10 phosphorylation *in cellulo*, as well as their effect on *in vitro* LRRK2 kinase activity toward a peptide and a Rab-GTPase substrate. This allows us to classify these Nbs in five functional groups (Table 1 & Fig. 6). Nbs belonging to functional group1 (containing Nb1, Nb6, Nb23) inhibit all tested LRRK2 kinase activities, including auto- and Rab phosphorylation in cells and peptide and Rab phosphorylation *in vitro*. This indicates that these Nbs probably block LRRK2 kinase activity *per se*. Group2 Nbs (containing Nb17, Nb36, Nb38 and potentially Nb40) only inhibited Rab phosphorylation in cells and *in vitro*, while leaving autophosphorylation and peptide phosphorylation unaffected. This pattern suggests that these Nbs probably either sterically interfere with binding of the Rab substrates or stabilize LRRK2 in a conformation that precludes Rab binding. Group3 Nbs (containing Nb22) activated LRRK2 kinase activity both in cells and *in vitro*. The group4 Nb, Nb42, had a strong inhibitory effect on LRRK2 autophosphorylation and Rab phosphorylation in cells, while leaving LRRK2 activity unaffected *in vitro*. At this moment, we can only speculate that this Nb specifically influences the in-cell behavior of LRRK2, potentially by influencing important protein-protein interactions or by preventing Rab29-mediated recruitment to a relevant sub-cellular compartment, such as the Golgi. Finally, binding of the fifth group of Nbs (Nb39 and potentially Nb40) does not influence LRRK2 kinase activity.

**Figure 6:**
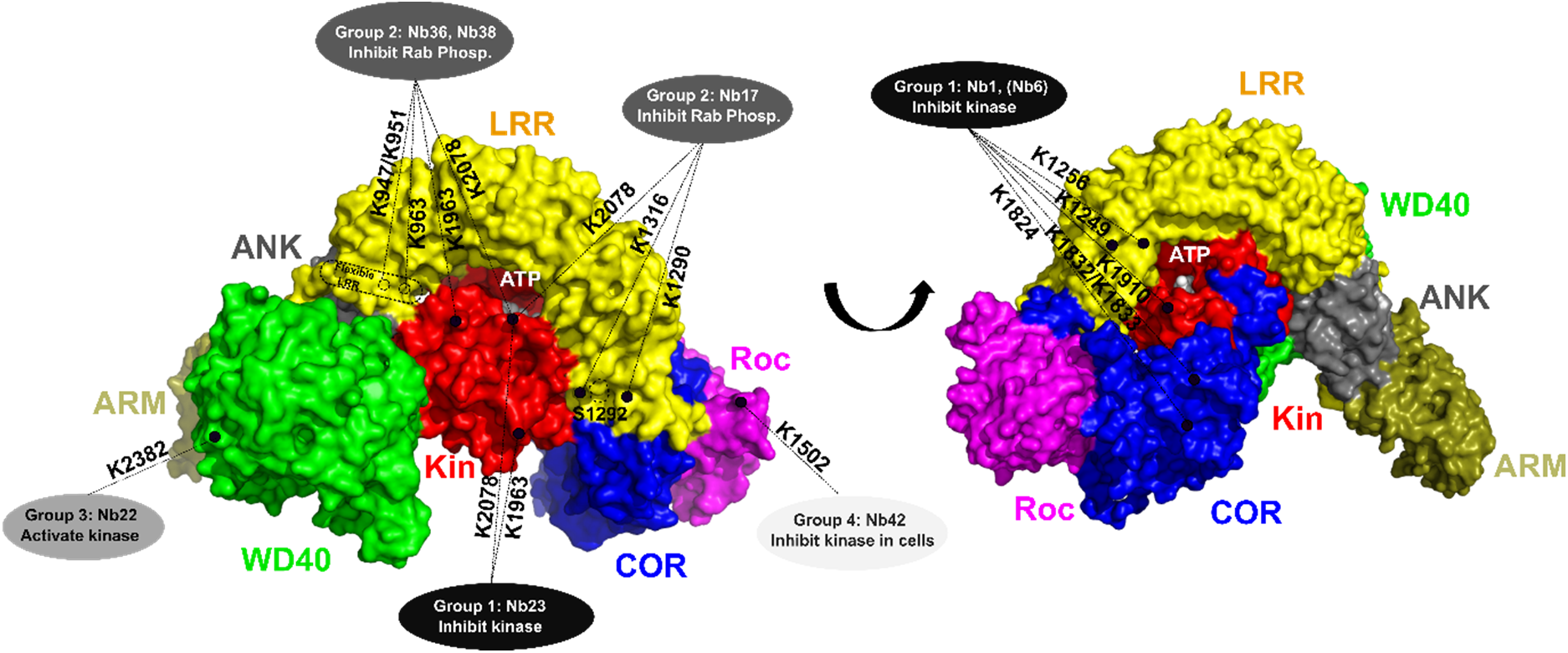
Schematic representation of the relation between the activity and binding epitopes of the different functional Nb groups. A surface representation of the cryo-EM structure of full-length LRRK2 is shown with the domains colored as indicated (PDB 7LHW) (29). The ATP binding pocket and the S1292 auto-phosphorylation site are indicated. The binding epitopes, determined by combining the results from ELISA and CL-MS experiments, of the Nbs belonging to different functional groups are indicated with dotted lines, with the LRRK2 lysine residues that form cross-links with the respective Nbs indicated adjacent to the lines.

The CL-MS experiments in combination with the recent cryo-EM structures of FL-LRRK2 and its C-terminal RCKW construct (27, 29) also allow us to map the approximate binding positions of the Nbs and to speculate on the mechanism of inhibition or activation of the group 1, 2 and 3 Nbs (Fig. 6, Fig. S13). The kinase inhibitory group1 Nbs fall in two categories based on their binding domains. According to ELISA-based domain mapping, Nb1 and Nb6 are able to bind exclusively to the C-terminal part of the COR domain (COR-B), and the CL-MS data for Nb1 suggest close contacts with residues of the COR-B, LRR and kinase domains (Fig. 2a, Fig. 6, Fig. S13a). An overall view on this cross-linking pattern mapped on the FL-LRRK2 structure (Fig. S13a) suggests that Nb1 binds via COR-B in a very central cavity in the middle of the interfaces formed by the LRR, COR-B, Roc and kinase domains. As such, we could hypothesize that the inhibition observed for Nb1 and Nb6 could be due to either sterically “pushing” LRRK2 toward an inactive more open conformation or “pulling” it together toward a closed inhibited conformation. The structure of FL-LRRK2 that is currently available indeed presumably shows LRRK2 in an inactive conformation where the LRR domain wraps around the kinase domain thereby keeping the kinase in an inactive conformation that is not accessible to substrates (29). Nb1 and Nb6 could act by stabilizing this inactive conformation. Another intriguing observation is that Nb1 also forms a cross-link with K1910 on the N-terminal lobe of the kinase domain. This latter residue is located between K1906 and E1920, two of the conserved regulatory triad residues formed by K1906-E1920-D2017. In the inactive state K1906 and E1920 interact with Y2018, thereby preventing a direct salt bridge between K1906 and E1920 and thus keeping the kinase in an inactive state (29). It has indeed been shown that via this mechanism Y2018 acts as a break on the kinase activity (38). Potentially, Nb1 can contribute to the stabilization of this inactive conformation by contacting the peptide region connecting K1906 and E1920 (Fig. S13a). On the other hand, the other group 1 Nb, Nb23, binds to the K-WD40 domain in ELISA and cross-links exclusively with K2078 and K2091 in the C-terminal lobe of the kinase domain (Fig6, Fig. S13b). The latter residues are located in relative close proximity to each other, but are located quite far from the ATP binding pocket. Interestingly, both cross-linking sites lay adjacent of residue N2081, of which the N2081D mutation has been identified as a major susceptibility factor for Crohn’s disease (6). N2081 makes a hydrogen bond with N1269 within the LRR domain, which might contribute to the stabilization of the LRR domain in an inactivating conformation, and, correspondingly, the N2081D mutation has been shown to increase Rab10 phosphorylation. We could hence speculate that Nb23 inhibits LRRK2 kinase activity by further locking LRRK2 in this inactive conformation. The observation that none of the group1 Nbs directly binds in the kinase ATP-binding pocket, while Nb1 and Nb6 even bind mainly outside the kinase domain, indicates that these Nbs do not act via an ATP-competitive mechanism, in agreement with our detailed kinetic analysis. Correspondingly, none of these Nbs induce microtubule relocalization of LRRK2 in contrast to ATP-competitive type 1 inhibitors.

All group2 Nbs (with the potential exception of Nb40) belong to what we called “full-length LRRK2 binders” that bind on the interface of N-terminal and C-terminal domains. The CL-MS data show that Nb17 binds close to K1290 and K1316 on the LRR domain and to K2078 in the C-terminal lobe of the kinase domain (Fig. 6, Fig.S13c). These three residues are in very close proximity to each other in the FL-LRRK2 structure, again in proximity to the “Crohn’s disease residue” N2081. The binding epitopes of Nb17 and Nb23 thus seem to partially overlap, although Nb23 is not interacting directly with the LRR domain (Fig. 6, Fig.S13c). Nb17 could hence inhibit the phosphorylation of bulky substrates, like Rab proteins, by keeping the LRR domain in the closed inhibitory conformation, while still a certain degree of flexibility or conformational change would be possible to allow the phosphorylation of smaller peptide substrates and autophosphorylation.

Nb36 and Nb38, which belong to the same Nb family, cross-link with residues on the C-terminal lobe of the kinase domain (K1963 and K2078) and the N-terminus of the LRR domain (K947, K951, K963). Although the latter three residues are not resolved in the cryo-EM structure of FL-LRRK2, this again suggest a mechanism similar to Nb17 where the LRR domain is blocked in an inhibitory conformation (Fig. 6). Together, these observations suggest a common mechanism for the group2 Nbs, which bind on the interface of several LRRK2 domains, thereby keeping LRRK2 in a “tight” conformation with the LRR domain wrapped as a lid around the kinase domains and occluding entry of bulky Rab substrates. In this respect it is also worth noting that “unleashing” of the N-terminal domains from the catalytic RCKW domains, as induced by type-1 kinase inhibitors and multiple PD mutations, has been linked to the association of LRRK2 to microtubules. Correspondingly, we found that group2 Nbs inhibit MLi-2-induced LRRK2 microtubule association (Fig. 5).

Finally, the kinase-activating Nb22 cross-links with residue K2382 on the WD40 domain. This binding epitope is very close to the interface of the WD40 domain with the ankyrin domain and the so-called “hinge-helix” of the LRR domain (aa 832-854). This hinge helix bridges the ARM, ANK and WD40 domains and thereby inhibits LRRK2 oligomerization via the WD40 domains, which in turn is associated with filament formation (29) (Fig 6, Fig. 13d). K2382 is located very close to G2385. The G2385R mutation is a PD associated variant that is common in the Chinese population (39) and was shown to disrupt WD40 dimer formation and the formation of pathogenic filaments in cells (28, 40). In very good agreement, we also find that Nb22 very efficiently inhibits MLi-2 filament formation in our experiments (Fig. 5). Moreover, probably via disrupting the interaction between the WD40 and the LRR hinge helix, the G2385R mutation leads to an activation of Rab10 phosphorylation (40), and we can thus anticipate that Nb22 increases the kinase activity via a similar mechanism.

The discovery of Nbs with a wide variety of LRRK2-modulating activities and with different mechanisms of inhibition forms a treasure chest for the further development of research tools to study the cellular role of LRRK2 and the associated disease mechanisms, as well as for future development of new diagnostic and therapeutic strategies. In particular, the finding that a subset of Nbs inhibit total LRRK2 kinase activity (group1), while others only inhibit Rab phosphorylation (group2), can be exploited to disentangle the role of the different LRRK2 activities and substrates in pathology. Owing to their ease of recombinant production, small size and stability, Nbs are also being intensively exploited for the development of diagnostics and therapeutics (23). This presents the Nbs described here as a potentially promising route for the development of new PD therapeutics next to other currently explored strategies, including but not restricted to small molecule kinase inhibitors (34, 41), inhibitors of GTP binding (42) and LRRK2-targetting antisense oligonucleotides (ASO) (43, 44). However, therapeutic targeting of LRRK2 has so far mainly focused on the development of small-molecule ATP-competitive inhibitors, and recently two molecules from Denali (DNL151 and DNL201) entered clinical trials (45, 46). But, despite these successes, some concerns regarding the observed adverse side effect upon prolonged treatment with high concentrations of ATP-competitive inhibitors remain (19, 20, 37). Recent studies identified oligomerization of overexpressed LRRK2 on microtubules, with concomitant blocking of microtubule-associated motor proteins, as a potential underlying cause of LRRK2 pathology. Similar to four out of the five common PD mutations, also treatment with type-1 ATP-competitive inhibitors increases such microtubule association. In contrast, the LRRK2 inhibiting Nbs act via completely novel allosteric mechanisms and inhibit LRRK2 activity via a mixed (non-competitive) type inhibition mechanism. Correspondingly, none of these Nbs induce LRRK2 association with microtubules, while some even revert MLi-2-induced LRRK2 relocalization. These observations will set the stage for further development of those Nbs into LRRK2 inhibitors with a completely different mode of action and cellular profile to the currently existing small molecule inhibitors. While Nbs have already been tested as therapeutics at the (pre-)clinical stage for different diseases, therapeutic targeting of LRRK2 with Nbs presents a number of challenges, including the need for obtaining sufficient levels of Nb within cells in the brain for a sustained period. One possible approach is to deliver the genes encoding the Nbs using a viral vector-based system (47). The field of gene therapy for the nervous system has undergone explosive growth in the last 5 years (48, 49), and gene therapy strategies have also been developed for PD (50–53). Different approaches for the development of the Nbs into next-generation therapeutics to target LRRK2 in PD are currently being explored.

## Materials and Methods

### Expression and purification of LRRK2 and LRRK2 constructs

Full-length human LRRK2 was expressed and purified based on previously developed protocols (26, 54), with minor adaptations to obtain purified LRRK2 either bound to Guanosine-5’-(γ-thio)- triphosphate (GTPγS) or Guanosine-5’-diphosphate (GDP) as described in the Supplementary Materials and Methods. Also the expression and purification of the LRRK2 4-domain Roc-COR-kinase-WD40 (RCKW) construct, the 2-domain kinase-WD40 (K-WD40) construct and the Roc-COR, Roc and COR-B domains are described in detail in the Supplementary Materials and Methods.

### Immunizations and Nb selection

In total three immunizations using different llamas were performed: (1) with the RocCOR domain construct of human LRRK2; (2) with full-length LRRK2 in the presence of an excess of GTPγS; and (3) with full-length LRRK2 in the presence of an excess of GDP. Additionally, in case of immunization 2 and 3, a mild crosslinking was performed on the protein after purification and prior to immunization in order to “trap” at least part of the injected proteins in their respective nucleotide-specific conformation during immunization (note that during all phage display selection steps non-crosslinked LRRK2 was used since a large excess of the nucleotides could be maintained in all these steps). For crosslinking, LRRK2 protein either loaded with 1 mM GTPγS or 1 mM GDP was incubated with the primary amine-specific crosslinker disuccinimidyl suberate (DSS) in 1:20 molar ratio for 30 minutes, after which the reaction was quenched by adding an excess of Tris.

A six-week protocol with weekly immunizations in presence of GERBU adjuvant was used, and blood was collected 4 days after the last injection. All animal vaccinations were performed in strict accordance with good practices and EU animal welfare legislation. Next, construction of immune libraries and Nb selection via phage display were performed using previously described protocols (55), as described in detail in the Supplementary Materials and Methods.

### Nanobody expression, purification and binding analysis via ELISA and pull-down experiments

The expression and purification of Nbs, as well as their analysis via ELISA and pull-down experiments were performed according to standard and previously published protocols (24) and are described in detail in the Supplementary Materials and Methods.

### *In cellulo* phospho-Rab assay

HEK293T cells cultured in DMEM supplemented with 10% Fetal Bovine Serum and 0.5% Pen/Strep, were transfected at a confluency of 50-70% with the individual Nb-GFP expression constructs, SF-tagged LRRK2(G2019S) and FLAG-HA Rab29. After 48 hrs cells were lysed and lysates were cleared by centrifugation and adjusted to a protein concentration of 1 µg/µl in 1x Laemmli Buffer. LRRK2 pS1292 and Rab10 T73 phosphorylation levels were determined by Western blot analysis as detailed in the Supplementary Materials and Methods. In brief, after SDS– PAGE separation, transfer to PVDF membranes and blocking, phospho-Rab10 levels were determined by the site-specific rabbit monoclonal antibody anti-pRAB10(pT73) (Abcam, ab230261), and LRRK2 autophosphorylation was determined by the site-specific rabbit monoclonal antibody anti-pLRRK2(pS1292) (Abcam, ab203181). Total LRRK2, Rab10 and Nb-GFP levels were determined by an in-house rat monoclonal antibody (clone 24D8) (56), the rabbit monoclonal antibody anti-RAB10/ERP13424 (Abcam, ab181367) and the rat monoclonal antibody anti-GFP (clone 3H9, ChromoTec), respectively. For detection, goat anti-rat IgG or anti-rabbit IgG HRP-coupled secondary antibodies (Jackson ImmunoResearch) were used.

### Chemical crosslinking/ mass spectrometry (CL-MS)

For chemical crosslinking, the LRRK2 protein solution was adjusted to a concentration of 3µM (0.86 mg/mL) and each Nb was added to LRRK2 at a final molar ratio of 2:1 and incubated for 1 h at 4 °C under constant mixing. The crosslinking reaction was then performed using the NHS-ester-based and CID-cleavable reagent disuccinimidyl sulfoxide (DSSO; Thermo Fisher) (57) at a molar excess of 60:1 (referred to the Nbs). The crosslinking reaction was carried out for 30 min at room temperature under constant mixing and stopped by adding Tris-HCl (pH 7.5) solution to a final concentration of 10 mM and incubation of 15 min at room temperature. Proteins were finally precipitated by chloroform/ methanol and subsequently subjected to tryptic proteolysis as described in (58). The tryptic peptide solutions were cleaned up by StageTips and subjected to SEC separation to enrich for crosslinked peptides as described earlier (26). Vacuum-dried fractions containing the crosslinked peptides, were analyzed individually on an Orbitrap Fusion mass spectrometer (Thermo Fisher) using the MS2_MS3 fragmentation method with the default settings (ver. 3.0, build 2041), and the Thermo Raw files were analyzed with the MS2_MS3 workflow provided by in Proteome Discoverer 2.5 (build 2.5.0.400), which uses XlinkX (ver. 2.5) (59) for the detection of crosslinked peptides, as described in detail in the Supplementary Material and Methods.

### Microscale thermophoresis and bio-layer interferometry measurements

Microscale thermophoresis (MST) experiments were performed for determining equilibrium binding affinities (K_D_) of Nbs binding to FL-LRRK2. Nbs were site-specifically labelled at their C-terminus with m-TAMRA, using Sortase-mediated peptide exchange (60). The Nbs were recloned into a pHEN29 vector which is subsequently used to express and purify the Nbs with a C-terminal LPETGG-His_6_-EPEA tag. Exchange of the latter peptide with an m-TAMRA-labeled GGGYK peptide (GenicBio, Shanghai, China) was performed using Sortase, and unlabeled Nb and unincorporated peptide were removed using a Ni-NTA and SEC purification step, respectively. MST measurements were performed using a Monolith NT.115 instrument (Nanotemper technologies) by titrating a fixed concentration of m-TAMRA-labelled Nb (50-100 nM) with varying concentrations of LRRK2 (16 points, 3:1 dilution series). Experiments were performed in 50 mM HEPES pH8.0, 150 mM NaCl, 10 mM MgCl_2_, 5% Glycerol, 0.1% BSA, 0.05% Tween and 500 µM GDP (except for Nb42 where 500 µM GTPγS was used). After incubation at 4°C for 30 mins, samples were loaded in capillaries and measurements were performed at 25°C using 50-70% LED power and 80% laser intensity (laser on-time: 30 s, laser off-time: 5 s). All experiments were performed in triplicate. Data was initially processed using the MO.affinity analysis software and final K_D_ values were obtained by fitting the MST signal (at 5-15 s on-time) *versus* [LRRK2] curves to a quadratic binding isotherm in GraphPad Prism 7.

Biolayer interferometry (BLI) measurements were performed using an Octet Red96 (FortéBio, Inc.) system in the same buffer, at 25 °C and with shaking speed 1000 rpm. Binding of the Nbs to the RCKW construct were performed using RCKW randomly biotinylated on lysine residues (bio-RCKW) loaded onto Streptavidin-coated (SA) biosensors at a concentration of 10 µg/mL. Association and dissociation kinetics were monitored for 600 s each, by first transferring the biosensor into increasing concentrations of Nb, followed by transferring them back to the wells containing buffers. Binding of Nbs to FL-LRRK2 were performed by first trapping LRRK2 on Streptavidin-coated (SA) biosensors by means of a high-affinity LRRK2-specific biotinylated Nb (either Nb40 or Nb42) loaded on SA biosensors at a concentration of 5 µg/mL. This Nb-loaded sensor was then used to trap FL-LRRK2 from a 50 nM LRRK2 solution. Finally, this sensor was used to monitor association (600 s to 1000 s) and dissociation (600 s to 1000 s) of the whole set of Nbs (to assess binding of Nb40 a sensor with Nb42 as trapping agent was used, for all other Nbs Nb40 was used as trapping agent). All experiments were performed in triplicate. The association / dissociation traces were fitted on a 1:1 binding model using either the local partial or global (full) options (implemented in the FortéBio Analysis Software). The resulting R_eq_ values were subsequently plotted against the Nb concentration, and used to derive the K_D_ values from the corresponding dose-response curves fitted on a Langmuir model. The fitted curves were exported, and final figures were generated using GraphPad Prism7.

### *In vitro* peptide phosphorylation assay

The LRRK2 kinase activity was determined using the PhosphoSens® Protein Kinase Assay (AssayQuant Technologies Inc.) using the optimized LRRK2 AQT0615 peptide as substrate and according to the manufacturers’ instructions. Each reaction mixture contained 10 µM AQT0615 peptide substrate/probe, 1 mM ATP and 500 µM of either GDP or GTPγS, in a buffer consisting of 50 mM HEPES pH 7.5, 0.1 % Brij-35, 50 mM NaCl and 10 mM MgCl_2_. The reaction was initiated by addition of a final concentration of 80 nM of LRRK2 either in absence or presence of 25 µM Nb (after pre-incubation of LRRK2 and Nbs for 30 minutes on ice). The LRRK2-catalyzed phosphorylation of the peptide substrate/probe is followed continuously at 30°C in a plate reader with excitation and emission wavelength of 360 nm and 485 nm respectively. Time traces were corrected by subtracting the “no LRRK2” control. The initial rates were determined from the slope of the linear portion of the curve.

To determine IC_50_ values the Nb concentration was varied from 150 µM to 0.006 µM using a two-fold serial dilution, at final concentrations of 150 nM LRRK2, 10 µM AQT0615 peptide, 1mM ATP and 500 µM GDP. The relative LRRK2 activity (compared to the “no nanobody” control) was plotted against the logarithmic Nb concentration and fitted on a three-parameter log(inhibitor) vs response equation, using GraphPad Prism 7. All time traces were collected in triplicate.

The mechanism of inhibition and apparent K_I_(K_I_^app^) values of Nb1, Nb6 and Nb23 were determined using the same assay. Full Michaelis-Menten curves in presence of different Nb concentrations were collected at 30°C in the same buffer as above (in presence of 500 mM GDP), using AQT0615 at 10 µM and at varying concentrations of ATP. The LRRK2 concentration was chosen such that initial rates (linear fluorescence versus time curves) were obtained, and LRRK2, ATP and the Nbs were preincubated at 4°C for 30min prior to starting the reaction by adding the peptide substrate. The Michaelis-Menten curves for the different Nb concentrations were globally fit on a mixed-inhibition model using GraphPad Prism 7, according to the equation.

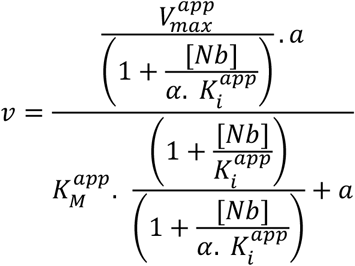

(with K_ic_^app^ = K_i_^app^ and K_iu_^app^ = α.K_i_ ^app^)

### *In vitro* Rab8a phosphorylation assay

To determine LRRK2 kinase activity toward Rab8a 100 nM LRRK2 and 25 µM of the respective Nbs were incubated for 30 min at 4°C in kinase assay buffer (50 mM HEPES pH 7.5, 50 mM NaCl, 10 mM MgCl_2_, 500 µM GDP or GTPγS, 1 mM ATP and 0.1 mg/ml BSA). A positive control, containing no Nb, and two negative controls, containing either 25 µM MLi-2 or no LRRK2, were treated the same way. The protein kinase reaction was initiated by adding 2.5 µM His6-Rab8a as a substrate and performed for 3 min at 30°C on an Eppendorf thermomixer at 650 rpm. To terminate the reaction, 5xSDS-sample buffer was added to the respective reactions and samples were subsequently incubated at 105°C for 10 min. For Western blotting, samples were transferred after SDS-PAGE via semi-dry blotting (Rab8a) or tank blotting (LRRK2) to 0.45µm nitrocellulose (GE Healthcare). Membranes were blocked with 5 % (w/V) BSA in TBS-T (Tris buffered saline supplemented with 0.1 % Tween 20) for 1 h. Monoclonal mouse-anti-His6 (1:1000, GE Healthcare) and monoclonal rabbit-anti-pT72-Rab8a (1:1000, Abcam) were used as primary antibodies and incubated at 4°C overnight. For LRRK2 detection, a monoclonal mouse-anti-Flag antibody (1:1000, SIGMA) was used as primary antibody. After washing, the membranes were incubated with secondary antibodies (IRDye®680RD Goat-anti-Mouse, IRDye®800CW Donkey-anti-Rabbit; both 1:15000, Li-COR) for 1 h and washed. Evaluation and quantification of the Western blots were performed by using an Odyssey FC imaging system and Image Studio Lite software version 5.2 (both LiCOR). The ratio of the pT72-Rab8a signal to the according total Rab8a signal was used for quantification. Additionally, the signal ratios were normalized by using the positive control without Nb as a reference.

For determination of the combined linear range of the anti-Rab8a pT72-antibody and anti-His6-antibody, a kinase assay, as described above, without Nb was performed for 1 h to achieve a full signal. An increasing amount of protein ranging from 4 – 500 ng was loaded on the SDS PAGE, blotted and quantified employing the Empiria Studio® 1.3 software (LiCOR). For subsequent experiments 20 ng Rab8a was loaded to the SDS-gel. To determine (semi-quantitative) dose-response curves, Nbs in concentrations ranging from 4 nM to 40 µM were incubated with 2.5 µM Rab8a as described above and Rab8a phosphorylation was quantified via WB-analysis (upon loading of 250 ng Rab8a on SDS-gel). Relative values were quantified with Image Studio Lite software version 5.2 (LiCOR) and dose response curves plotted against the concentration of the respective Nbs were calculated using GraphPad Prism 6.

### Confocal microscopy and microtubule localization

A HEK 293 cell line was cultured in complete media (high-glucose Dulbecco’s modified Eagle’s medium, 10% fetal bovine serum and Penicillin-Streptomycin-Glutamine [Gibco]). Cells were seeded on 8-well µ-Slide(Ibidi) and transfected at a confluency of 50-70% with GFP-Nb and mScarlet-LRRK2 constructs, using JetPEI reagent (Polyplus transfection). After 24 hrs, cells were treated with either DMSO or 1 µM of LRRK2 kinase inhibitor (MLi-2, cat.no. 5756, TOCRIS) for 90 mins and then examined for localization. Data acquisition was done with a ×100 oil-immersion objective with a Zeiss LSM800 confocal laser scanning microscope. Image analysis of z-scan was done using the Zeiss microscope software ZEN.

## Acknowledgements

This work was supported by the Michael J. Fox Foundation for Parkinson’s Research (grant number 14527 to A.K., C.J.G., W.V., and 14527.01 to F.W.H., A.K., C.J.G., W.V.), the Fonds voor Wetenschappelijk Onderzoek (G005219N to W.V) and a Strategic Research Program Financing from the VUB (SRP50 to W.V.). L.V.R. was supported by a VUB OZR bridging grant. We acknowledge Instruct-ERIC, part of ESFRI, and the FWO for their support to the Nanobody discovery. The authors thank the staff of the Core Facility for Medical Bioanalytics at the Institute for Ophthalmic Research (University of Tubingen) for technical assistance and Bernd Gilsbach for help in LRRK2 protein purification. We thank Sebastian Mathea for fruitful scientific discussions.

## Author Contributions

R.K.S. purified proteins, generated the Nbs and performed most *in vitro* biochemical/biophysical experiments, with the help of L.V.R. and T.D.M.; G.G. produced proteins and performed the XL-MS experiment; A.S. and F.v.Z performed most cell-based assays; E.S. and S.H.S. performed the *in vitro* Rab kinase assays; D.C. and S.K. produced and purified proteins; E.P. and J.S. supervised the immunization, Nb library construction and Nb discovery; E.J.K., F.W.H., A.K., C.J.G. and W.V. conceived, supervised and interpreted experiments. A.K., C.J.G and W.V. conceived the study. R.K.S. and W.V. wrote the manuscript with the help and input of all other authors.

## Materials Availability Statement

All Nanobodies that are characterized in this study are available from the Versées Lab upon request by contacting.

## References

1. A. Elbaz, L. Carcaillon, S. Kab, F. Moisan, Epidemiology of Parkinson’s disease. Rev Neurol 172, 14–26 (2016).

2. E. Monfrini, A. Di Fonzo, “Leucine-rich repeat kinase (LRRK2) genetics and parkinson’s disease” in Advances in Neurobiology, (2017), pp. 3–30.

3. R. Di Maio, et al., LRRK2 activation in idiopathic Parkinson’s disease. Sci. Transl. Med. 10, eaar5429 (2018).

4. W. P. Gilks, et al., A common LRRK2 mutation in idiopathic Parkinson’s disease. Lancet 365, 415–416 (2005).

5. J. C. Barrett, et al., Genome-wide association defines more than 30 distinct susceptibility loci for Crohn’s disease. Nat. Genet. 40, 955–62 (2008).

6. K. Y. Hui, et al., Functional variants in the LRRK2 gene confer shared effects on risk for Crohn’s disease and Parkinson’s disease. Sci.Transl.Med. 10, eaai7795 (2018).

7. L. Wauters, W. Versées, A. Kortholt, Roco Proteins: GTPases with a Baroque Structure and Mechanism. Int J Mol Sci. 20, 147 (2019).

8. M. Steger, et al., Phosphoproteomics reveals that Parkinson’s disease kinase LRRK2 regulates a subset of Rab GTPases. Elife 5, e12813. (2016).

9. Z. Liu, et al., LRRK2 phosphorylates membrane-bound Rabs and is activated by GTP-bound Rab7L1 to promote recruitment to the trans-Golgi network. Hum. Mol. Genet. 27, 385–395 (2017).

10. A. Marchand, M. Drouyer, A. Sarchione, M. C. Chartier-Harlin, J. M. Taymans, LRRK2 Phosphorylation, More Than an Epiphenomenon. Front. Neurosci. 14, 527 (2020).

11. E. Deyaert, et al., A homologue of the Parkinson’s disease-associated protein LRRK2 undergoes a monomer-dimer transition during GTP turnover. Nat. Commun. 8, 1008 (2017).

12. S. Sen, P. J. Webber, A. B. West, Dependence of leucine-rich repeat kinase 2 (LRRK2) kinase activity on dimerization. J. Biol. Chem. 284, 36346–36356 (2009).

13. Z. Berger, K. A. Smith, M. J. Lavoie, Membrane localization of LRRK2 is associated with increased formation of the highly active lrrk2 dimer and changes in its phosphorylation. Biochemistry 49, 5511–5523 (2010).

14. P. A. Lewis, et al., The R1441C mutation of LRRK2 disrupts GTP hydrolysis. Biochem. Biophys. Res. Commun. 357, 668–671 (2007).

15. A. B. West, et al., Parkinson’s disease-associated mutations in leucine-rich repeat kinase 2 augment kinase activity. PNAS 102, 16842–16847 (2005).

16. Z. Sheng, et al., Ser1292 autophosphorylation is an indicator of LRRK2 kinase activity and contributes to the cellular effects of PD mutations. Sci. Transl. Med. 4, 164ra161 (2012).

17. J.-M. Taymans, E. Greggio, LRRK2 Kinase Inhibition as a Therapeutic Strategy for Parkinson’s Disease, Where Do We Stand? Curr. Neuropharmacol. 14, 214–25 (2016).

18. S. Domingos, T. Duarte, L. Saraiva, R. C. Guedes, R. Moreira, Targeting leucine-rich repeat kinase 2 (LRRK2) for the treatment of Parkinson’s disease. Future Med. Chem. 11, 1953–1977 (2019).

19. R. N. Fuji, et al., Effect of selective LRRK2 kinase inhibition on nonhuman primate lung. Sci Transl Med. 7, 1–13 (2015).

20. M. A. S. Baptista, et al., Loss of leucine-rich repeat kinase 2 (LRRK2) in rats leads to progressive abnormal phenotypes in peripheral organs. PLoS One 8, e80705 (2013).

21. D. Bryce, et al., Characterization of the onset, progression, and reversibility of morphological changes in mouse lung following pharmacological inhibition of LRRK2 kinase activity. J. Pharmacol. Exp. Ther., Online ahead of publication. (2021).

22. X. Lu, J. B. Smaill, K. Ding, New Promise and Opportunities for Allosteric Kinase Inhibitors. Angew Chem Int Ed Engl. 59, 13764–13776 (2019).

23. S. Muyldermans, Nanobodies: Natural Single-Domain Antibodies. Annu. Rev. Biochem. 82, 775–797 (2013).

24. M. Leemans, et al., Allosteric modulation of the GTPase activity of a bacterial LRRK2 homolog by conformation-specific Nanobodies. Biochem J. 477, 1203–1218 (2020).

25. J. H. Kluss, et al., Detection of endogenous S1292 LRRK2 autophosphorylation in mouse tissue as a readout for kinase activity. npj Park. Dis. 4 (2018).

26. G. Guaitoli, et al., Structural model of the dimeric Parkinson’s protein LRRK2 reveals a compact architecture involving distant interdomain contacts. Proc. Natl. Acad. Sci. 113, E4357–E4366 (2016).

27. C. K. Deniston, et al., Structure of LRRK2 in Parkinson’s disease and model for microtubule interaction. Nature 588, 344–349 (2020).

28. R. Watanabe, et al., The in situ structure of Parkinson’s disease-linked LRRK2. Cell 82, 1508–1518 (2020).

29. A. Myasnikov, et al., Structural analysis of the full-length human LRRK2. Cell 184, 1–9 (2021).

30. A. Gardet, et al., LRRK2 Is Involved in the IFN-γ Response and Host Response to Pathogens. J. Immunol. 185, 5577–85 (2010).

31. J. D. Scott, et al., Discovery of a 3-(4-Pyrimidinyl) Indazole (MLi-2), an Orally Available and Selective Leucine-Rich Repeat Kinase 2 (LRRK2) Inhibitor that Reduces Brain Kinase Activity. J. Med. Chem. 60, 2983–2992 (2017).

32. M. J. Fell, et al., MLi-2, a potent, selective, and centrally active compound for exploring the therapeutic potential and safety of LRRK2 kinase inhibition. J. Pharmacol. Exp. Ther. 355, 397–409 (2015).

33. D. S. Williamson, et al., Design of Leucine-Rich Repeat Kinase 2 (LRRK2) Inhibitors Using a Crystallographic Surrogate Derived from Checkpoint Kinase 1 (CHK1). J. Med. Chem. 60, 8945–8962 (2017).

34. X. Ding, F. Ren, Leucine-rich repeat kinase 2 inhibitors: a patent review (2014-present). Expert Opin. Ther. Pat. 30, 275–286 (2020).

35. N. Dzamko, et al., Inhibition of LRRK2 kinase activity leads to dephosphorylation of Ser 910/Ser935, disruption of 14-3-3 binding and altered cytoplasmic localization. Biochem. J. 430, 405–413 (2010).

36. L. R. Kett, et al., LRRK2 Parkinson disease mutations enhance its microtubule association. Hum. Mol. Genet. 21, 890–899 (2012).

37. M. A. Andersen, et al., PFE-360-induced LRRK2 inhibition induces reversible, non-adverse renal changes in rats. Toxicology 395, 15–22 (2018).

38. S. H. Schmidt, et al., The dynamic switch mechanism that leads to activation of LRRK2 is embedded in the DFGψ motif in the kinase domain. Proc Natl Acad Sci USA 116, 14979–14988 (2019).

39. I. F. Mata, et al., Lrrk2 pathogenic substitutions in Parkinson’s disease. Neurogenetics 6, 171–177 (2005).

40. P. Zhang, et al., Crystal structure of the WD40 domain dimer of LRRK2. Proc. Natl. Acad. Sci. U. S. A. 116, 1579–1584 (2019).

41. M. A. S. Baptista, et al., LRRK2 inhibitors induce reversible changes in nonhuman primate lungs without measurable pulmonary deficits. Sci. Transl. Med. 12, eaav0820 (2020).

42. T. Li, et al., Novel LRRK2 GTP-binding inhibitors reduced degeneration in Parkinson’s disease cell and mouse models. Hum. Mol. Genet. 23, 6212–6222 (2014).

43. J. A. Korecka, et al., Splice-Switching Antisense Oligonucleotides Reduce LRRK2 Kinase Activity in Human LRRK2 Transgenic Mice. Mol. Ther. - Nucleic Acids 21, 623–635 (2020).

44. H. T. Zhao, et al., LRRK2 Antisense Oligonucleotides Ameliorate α-Synuclein Inclusion Formation in a Parkinson’s Disease Mouse Model. Mol Ther Nucleic Acids 8, 508–519 (2017).

45. Study to Evaluate DNL201 in Subjects With Parkinson’s Disease.

46. Study to Evaluate DNL151 in Subjects With Parkinson’s Disease.

47. A. Messer, D. C. Butler, Optimizing intracellular antibodies (intrabodies/nanobodies) to treat neurodegenerative disorders. Neurobiol. Dis. 134 (2020).

48. E. Hudry, L. H. Vandenberghe, Therapeutic AAV Gene Transfer to the Nervous System: A Clinical Reality. Neuron 101, 839–862 (2019).

49. S. Kimura, H. Harashima, Current status and challenges associated with CNS-targeted gene delivery across the BBB. Pharmaceutics 12, 1–33 (2020).

50. A. Merola, et al., Gene Therapy in Movement Disorders: A Systematic Review of Ongoing and Completed Clinical Trials. Front. Neurol. 12 (2021).

51. P. C. Buttery, R. A. Barker, Gene and Cell-Based Therapies for Parkinson’s Disease: Where Are We? Neurotherapeutics 17, 1539–1562 (2020).

52. F. L. Hitti, A. I. Yang, P. Gonzalez-Alegre, G. H. Baltuch, Human gene therapy approaches for the treatment of Parkinson’s disease: An overview of current and completed clinical trials. Park. Relat. Disord. 66, 16–24 (2019).

53. M. Hocquemiller, L. Giersch, M. Audrain, S. Parker, N. Cartier, Adeno-Associated Virus-Based Gene Therapy for CNS Diseases. Hum. Gene Ther. 27, 478–96 (2016).

54. C. J. Gloeckner, et al., Phosphopeptide analysis reveals two discrete clusters of phosphorylation in the N-terminus and the Roc domain of the Parkinson-disease associated protein kinase LRRK2. J. Proteome Res. 9, 1738–45 (2010).

55. E. Pardon, et al., A general protocol for the generation of Nanobodies for structural biology. Nat. Protoc. 9, 674–693 (2014).

56. M. D. P. Carrion, et al., The LRRK2 G2385R variant is a partial loss-of-function mutation that affects synaptic vesicle trafficking through altered protein interactions. Sci. Rep. 7, 5377 (2017).

57. W. V. Kandur, A. Kao, D. Vellucci, L. Huang, S. D. Rychnovsky, Design of CID-cleavable protein cross-linkers: Identical mass modifications for simpler sequence analysis. Org. Biomol. Chem. 13, 9793–807 (2015).

58. C. J. Gloeckner, K. Boldt, M. Ueffing, Strep/FLAG tandem affinity purification (SF-TAP) to study protein interactions. Curr. Protoc. Protein Sci. 57, 19.20.1-19.20.19 (2009).

59. F. Liu, D. T. S. Rijkers, H. Post, A. J. R. Heck, Proteome-wide profiling of protein assemblies by cross-linking mass spectrometry. Nat. Methods 12, 1179–84 (2015).

60. S. Massa, et al., Sortase A-mediated site-specific labeling of camelid single-domain antibody-fragments: a versatile strategy for multiple molecular imaging modalities. Contrast Media Mol. Imaging 11, 328–339 (2016).

