## Supplementary Information for "Nanobodies as allosteric modulators of Parkinson’s disease-associated LRRK2"

### Supplementary Materials and Methods

#### Expression and purification of LRRK2 and LRRK2 constructs

Full-length human LRRK2 was expressed and purified based on previously developed protocols (1, 2), with minor adaptations to obtain purified LRRK2 either bound to Guanosine-5'-(γ-thio)-triphosphate (GTPγS) or Guanosine-5'-diphosphate (GDP). Briefly, full-length LRRK2 was cloned into the N-Strep/Flag-TAP (N-SF-TAP) pcDNA3.0 vector coding for a protein with an N-terminal Twin-Strep-tag and FLAG-tag (NSF) (3). HEK293T cells (CRL-11268; American Type Culture Collection) were transfected between 50% and 70% confluence with 8 μg of plasmid DNA/ 14cm culture dish using polyethyleneimine (PEI) 25 kDa (Polysciences), and cultured post-transfection for 48 hrs in 14-cm dishes in DMEM (Sigma-Aldrich) supplemented with 10% (vol/vol) FBS (Sigma-Aldrich) and appropriate antibiotics. After removal of the medium, the cells were resuspended in lysis buffer (1 mL / 14-cm dish) containing 50 mM HEPES (pH 8.0), 150 mM NaCl, 5 mM $MgCl_2$, 2 mM DTT, 5% Glycerol and supplemented with 0.55% (v/v) Nonidet P-40, Complete™ EDTA-free Protease Inhibitor Cocktail (Roche) and either 1mM GTPγS or 1 mM GDP. After cell lysis, the lysate was incubated with Strep-Tactin beads (IBA), and after extensive washing with 50 mM HEPES (pH 8.0), 150 mM NaCl, 2 mM DTT, 5 mM $MgCl_2$, 0.5 mM EDTA, 10% [vol/vol] glycerol, containing either 1 mM GTPγS or GDP, elution was performed with the same buffer containing 2.5 mM of D-Desthiobiotin (IBA) and 1 mM of either GTPγS or GDP.

The LRRK2 4-domain Roc-COR-kinase-WD40 (RCKW) construct and the 2-domain kinase-WD40 (K-WD40) construct were expressed and purified according to a previously established protocol. Exponentially growing TriEx Sf9 insect cells (Novagen) were diluted to a density of $2x10^7$ cells/mL. Then, a high-titer virus suspension was added in the ratio 1:64. The viruses had been generated using the Bac-to-Bac expression system (Invitrogen) and the expression vector pFB-6HZB (SGC). The expressed proteins were composed of an N-terminal $His_6$-Z tag and the LRRK2 residues 1327-2527 (RCKW), and 1847-2527 (K-WD40), respectively. The infected cells were incubated in 800 mL aliquots in shaker flasks (66 h, gentle agitation, 27°C), harvested by centrifugation and stored at -20°C. For purification, the Sf9 cell pellets were washed with PBS, re-suspended in lysis buffer (50 mM HEPES pH 7.4, 500 mM NaCl, 20 mM imidazole, 0.5 mM TCEP, 5% glycerol, 5 mM $MgCl_2$, 20 μM GDP) and lysed by sonication. The lysates were cleared by centrifugation and loaded onto a Ni-NTA column. After vigorous rinsing with lysis buffer, the $His_6$-Z-tagged proteins were eluted in lysis buffer containing 300 mM imidazole. Immediately thereafter, the eluates were diluted with a buffer containing no NaCl, in order to reduce the NaCl-concentration to 250 mM and loaded onto an SP sepharose column. The $His_6$-Z-tagged proteins were eluted with a 250 mM to 2.5 M NaCl gradient and treated with TEV protease overnight to cleave the $His_6$-Z tag. Contaminating proteins, the cleaved tag, uncleaved LRRK2 protein and TEV protease were removed in another combined SP Sepharose Ni-NTA step. Finally, the LRRK2 constructs were concentrated and subjected to gel filtration in storage buffer (20 mM HEPES pH 7.4, 800 mM NaCl, 0.5 mM TCEP, 5% glycerol, 2.5 mM $MgCl_2$, 20 μM GDP) using an AKTA Xpress system combined with an S200 gel filtration column.

A domain construct of LRRK2 spanning the Roc and COR domains (RocCOR) was expressed from a pBADcLIC vector coding for a protein spanning the a.a. residues 1334-1840 fused to an N-terminal Twin-Strep-tag and C-terminal $His_{10}$-tag. The pBADcLIC vector containing the RocCOR coding region was transformed into an *E. coli* strain (*E. coli RCEv9*) that was custom evolved in-house starting from a MC1061Δ*acrB* strain for optimal expression of the RocCOR protein, using previously described protocols (4, 5). Protein expression was induced at an OD of about 0.7 with 0.01% of arabinose and allowed to proceed overnight at 20°C. Cells were resuspended and lysed in a buffer containing 50 mM Tris-HCl pH 7.5, 500 mM NaCl, 10 mM $MgCl_2$, 2mM β-mercaptoethanol, 20 mM imidazole and supplemented with 1 mM of PMSF, 1 μg/mL of Leupeptin, 0.1 μg/mL of AEBSF, 50 μg/mL of DNaseI and either 0.5 mM GDP or 0.5 mM Guanosine-5'-[( β,γ )-imido]triphosphate (GppNHp). The RocCOR protein was loaded on a Ni-NTA column, and after extensive washing with the same buffer, supplemented with 300mM KCl and 5mM ATP to reduce contamination with chaperones, the protein was eluted using the same buffer with 300mM imidazole. A final purification step consisted of a gel filtration on a Superdex S200 10/300 column using 30 mM HEPES pH 7.5, 150 mM NaCl, 5 mM $MgCl_2$, 5% glycerol, 1 mM DTT as a buffer supplemented with either 0.5 mM GDP or GppNHp.

A construct of the Roc domain spanning the residues 1329-1520 was cloned in the pET-28a vector providing an N-terminal $His_6$-tag, and the vector was transformed in the *E. coli* BL21(DE3) strain. At an OD of about 0.7 protein expression was induced with 0.1 mM isopropyl β-D-1-thiogalactopyranoside (IPTG) and allowed to proceed overnight at 20°C. Cells were resuspended and lysed into a buffer containing 30 mM HEPES pH 7.5, 250 mM NaCl, 10 mM $MgCl_2$, 10 mM glycine and 20 mM imidazole and supplemented with 1 mM of PMSF, 1 μg/mL of Leupeptin, 0.1 μg/mL of AEBSF and 50 μg/mL of DNaseI. Recombinant protein was purified using a two-step protocol consisting of a Ni-NTA affinity chromatography step followed by gel filtration on a Superdex S75 10/300 column using 30 mM HEPES pH 7.5, 150 mM NaCl, 5 mM $MgCl_2$, 5% glycerol, 1 mM DTT as buffer.

A construct of the C-terminal part of the COR domain (COR-B) spanning the residues 1672-1840 was cloned in the pDEST-566 vector providing an N-terminal $His_6$-MBP-tag, and the vector was transformed in the *E. coli* BL21(DE3) strain. When an OD of about 0.7 was reached protein expression was induced with 0.5 mM IPTG and allowed to proceed for 2h at 20°C. Cells were resuspended and lysed in a buffer containing 30 mM HEPES pH 7.5, 200 mM NaCl, 1mM EDTA and 1mM DTT and supplemented with 1 mM of PMSF, 1 μg/mL of Leupeptin, 0.1 μg/mL of AEBSF and 50 μg/mL of DNaseI. The cell lysate was loaded on a 5 mL MBPTrap column (GE Healthcare), and after washing with resuspension buffer, the protein was eluted in the same buffer containing 10 mM maltose. A final purification step consisted of a gel filtration on a Superdex S200 10/300 column using 30 mM HEPES pH 7.5, 150 mM NaCl as buffer.

**Immunizations and Nb selection**

In total three immunizations using different llamas were performed: (1) with the RocCOR domain construct of human LRRK2; (2) with full-length LRRK2 in the presence of an excess of GTPγS; and (3) with full-length LRRK2 in the presence of an excess of GDP. Additionally, in case of immunization 2 and 3, a mild crosslinking was performed on the protein after purification and prior to immunization in order to "trap" at least part of the injected proteins in their respective nucleotide-specific conformation during immunization (note that during all phage display selection steps non-crosslinked LRRK2 was used since a large excess of the nucleotides could be maintained in all these steps). For crosslinking,

LRRK2 protein either loaded with 1 mM GTPγS or 1 mM GDP was incubated with the primary amine-specific crosslinker disuccinimidyl suberate (DSS) in 1:20 molar ratio for 30 minutes, after which the reaction was quenched by adding an excess of Tris. A six-week protocol with weekly immunizations in presence of GERBU adjuvant was used, and blood was collected 4 days after the last injection. All animal vaccinations were performed in strict accordance with good practices and EU animal welfare legislation.

Next, construction of immune libraries and Nb selection via phage display were performed using previously described protocols (6). In brief, starting from the blood collected from the llamas after immunization, the variable domains of the heavy-chain antibody repertoire were cloned in a pMESy4 phage display vector, which adds a C-terminal $His_6$-tag and EPEA-tag (= CaptureSelect™ C-tag) upon protein expression. This resulted in three independent immune libraries of $8.3\times10^8$, $1.8\times10^9$ and $1.3\times10^9$ transformants, respectively. This Nb repertoire was expressed on the tip of filamentous phages after rescue with the VCSM13 helper phage. For all libraries, two consecutive rounds of phage display selection were performed against either solid phase coated full-length LRRK2 or full-length LRRK2 trapped on beads via anti-flag M2 Ab (Merck). Coating of LRRK2 was performed in a coating buffer containing 50 mM HEPES pH 8.0, 150 mM NaCl, 5 mM $MgCl_2$, 5% glycerol, while for all the binding and washing steps additionally 0.05 % Tween20 was added. Moreover, all buffers were supplemented with either 1mM GDP (immunization 1 and 3) or GTPγS (immunization 2) during all coating steps, and 100 μM of the same nucleotides during binding and washing steps. For the libraries resulting from immunization 2 and 3, additionally, three consecutive rounds of phage display selection were performed using the solid phase coated LRRK2 Roc domain. Several single colonies were picked after each round of phage display selection and sequence analysis was used to classify the resulting Nb clones in sequence families based on their CDR3 sequence.

**Nanobody expression and purification**

Nb expression and purification was performed as described previously (6). The Nb-coding open reading frames, cloned in the pMESy4 vector (GenBank KF415192), were transformed in non-suppressor *E. coli* WK6 ($Su^-$) cells. Cells were grown at 37°C in Terrific Broth (TB) medium and protein expression was induced with 1 mM IPTG. After overnight expression at 28°C, cells were harvested via centrifugation and subjected to an osmotic shock to obtain the periplasmic extract. Subsequently, an affinity purification step on $Ni^{2+}$-NTA Sepharose followed by a dialysis step against a buffer consisting of 20 mM Tris-HCl pH 7.5, 150 mM NaCl was used to purify the Nbs.

In addition, to allow expression of the Nbs as fluorescently (eGFP)-labeled intrabodies (Fluobodies) in HEK293(T) cells (7), the Nb ORFs were recloned to the pEGFP-N1 vector using the HindIII and BamHI restriction sites. This will result in expression of the Nbs fused at their C-terminus to eGFP (Nb-GFP). To allow expression of the Nbs in HEK293(T) cells without GFP, a stop codon was introduced between the Nb- and GFP-coding sequence.

**Enzyme-linked immunosorbent (ELISA) assays**

Before Nb purification, binding of the Nbs to LRRK2 was confirmed using an ELISA screen on crude extracts of *E. coli* cells expressing the respective Nbs. Full-length LRRK2 was solid phase-coated on the bottom of the ELISA well, and the coating, blocking, binding and washing buffers were kept the same as in the phage display experiments and were supplemented with the relevant nucleotides (GTPγS or GDP). Binding of the Nbs to LRRK2 was detected via their EPEA-tag using a 1:4000 CaptureSelect™ Biotin anti-C-tag conjugate (Thermo Fischer Scientific) in combination with 1:1000 Streptavidin Alkaline Phosphatase (Promega). Color was developed by adding 100 µl of a 4 mg/mL disodium 4-nitrophenyl phosphate solution (DNPP, Sigma-Aldrich) and measured at 405 nm.

After purification of the selected Nbs as described above, ELISA experiments were performed to determine their domain specificity. Full-length LRRK2 and the Roc, COR-B, RocCOR and K-WD40 domain constructs were solid phase coated in 96-well ELISA plates. The coating, binding and washing buffer were kept the same as in the phage display experiments, supplemented with 100 µM GDP. The ELISA was developed in the same way as described above.

To assess any potential competition between the binding of Nanobodies Nb1, Nb6 or Nb23 and Mli-2 a competition ELISA experiment was performed. The ELISA experiment was performed similar as described above, using a fixed concentration of LRRK2 coated on the bottom of the ELISA plate, and using a 1:4 dilution series of Nb1, Nb6 or Nb23 ranging from 450 nM to 0.2 nM, resulting in a dose response titration curve. This setup were performed in the absence or in the presence of either a large excess of MLi-2 (1 μM), or an excess of the corresponding non-tagged Nb (9μm). A "no antigen control", where no LRRK2 was coated on the bottom of the well, was also included. Measurements in presence of MLi-2 were done in triplicate.

**Pull-down experiments**

Fresh lysate of HEK293 cells overexpressing SF-tagged LRRK2 and GFP-tagged Nbs was prepared in 100 mL ice-cold lysis buffer (10 mM Tris/HCl pH 7.5, 150 mM NaCl, 0.5 mM EDTA, 0.5% NP-40), containing complete EDTA-free protease inhibitor cocktail (Sigma–Aldrich Cat # 11836170001) and Protease Inhibitor Cocktail (Sigma, cat. no. P-2714). GFP-Nbs were immunoprecipitated with Magnetic GFP nanotrap beads (ChromoTek)(8, 9). Immune complexes were washed twice with 10 mM Tris/HCl pH 7.5 and subjected to immunoblot analysis by boiling samples in sample buffer with a reducing agent. Samples were separated on 4-15 % Tris-Glycine gels (Mini-PROTEAN® TGX™ Precast Gels, Bio-rad), transferred onto a nitrocellulose membrane (GE Lifesciences), and processed for Western blot analysis. Membranes were blocked in 5% dry milk in Tris-buffered saline plus Tween-20 for 1 hour. To allow for separate detection of LRRK2 and Nbs, the membrane was cut horizontally at 180 kDa and the upper part was probed with rat monoclonal anti-LRRK2 (clone 24D8 1:1000, Gloeckner lab), while the lower part was probed with Rabbit anti-GFP antibodies, 1:2500 (MA5-15256, Invitrogen) and incubated overnight at 4 C with gentle shaking. Membranes were then washed three times for 10 min at room temperature in PBS containing 0.1% or 0.05% Tween-20 and then incubated for 1 hour with anti-rat IgG-HRP (sc-2750, Santa Cruz Biotechnology) for LRRK2 and anti-rabbit HRP conjugated (#7074, Cell Signaling, 1:5000) for GFP-Nbs. Membranes were again washed three times for 10 min at room temperature in PBS containing 0.1% or 0.05 Tween-20. The membranes were coated with enhanced chemiluminescent (ECL) reagent (WesternSure PREMIUM, Li-COR biosciences), and proteins were detected using the C-Digit Imaging System (Li-COR Biosciences)

To test for binding of Nbs with endogenous (mouse) LRRK2, fresh lysates of RAW264.7 cells were prepared in 10 mM Tris/HCl pH 7.5; 150 mM NaCl; 0.5% NP-40 supplemented with complete EDTA-free protease inhibitor cocktail (Sigma–Aldrich Cat # 11836170001) and Protease Inhibitor Cocktail (Sigma, cat. no. P-2714). Purified His-tagged Nbs were added to the lysate at a final concentration of 1.5 μM and the mixture was allowed to rotate at 4°C overnight. His-tagged Nbs were pulled down by magnetic Dynabeads (Invitrogen). Immune complexes were washed twice with 10 mM Tris/HCl pH 7.5 and subjected to immunoblot analysis in a similar way as described above. To allow for separate detection of LRRK2 and Nbs, the membrane was cut horizontally at 180 kDa and the upper part was probed with rabbit monoclonal anti-LRRK2, 1:1000 ([MJFF2 (c41-2)] , ab133474, Abcam) while the lower part was probed with mouse anti-Histidine Tag antibody, 1:1000 (AD1.1.10, Bio-rad) and incubated overnight at 4 °C with gentle shaking. Membranes were then washed three times for 10 min at room temperature in PBS containing 0.1% or 0.05% Tween 20 and then incubated for at least 1hour (light protected) with secondary antibodies: anti-rabbit HRP-conjugated (#7074, Cell Signaling, 1:500) or mouse IgG kappa binding protein (m-IgGk BP) conjugated to HRP ( sc-516102, Santa Cruz Biotechnology, 1:5000). Membranes were again washed three times for 10 min at room temperature in PBS containing 0.1% or 0.05% Tween-20. The membranes were coated with enhanced chemiluminescent (ECL) reagent (WesternSure PREMIUM, Li-COR biosciences), and proteins were detected by C-Digit Imaging System (Li-COR Biosciences).

**Western blot analysis of *in cellulo* LRRK2 kinase activity**

HEK293T cells were cultured in DMEM (supplemented with 10% Fetal Bovine Serum, 0.5% Pen/Strep). For the assay, the cells were seeded onto six-well plates and transfected at a confluency of 50-70% with the individual Nb-GFP expression constructs, SF-tagged LRRK2(G2019S) and FLAG-HA Rab29 using a self-made polyethylenimine (PEI)-based transfection reagent (10). After 48 hrs cells were lysed in lysis buffer [30 mM Tris-HCL (pH7.4), 150 mM NaCl, 0.5% Nonident-P40, complete protease inhibitor cocktail, phosphatase inhibitor cocktail II & III (all Sigma)]. Lysates were cleared by centrifugation at 10,000 x g and adjusted to a protein concentration of 1 μg/μl in 1x Laemmli Buffer. Samples were subsequently subjected to SDS PAGE and Western Blot analysis to determine LRRK2 pS1292 and Rab10 T73 phosphorylation levels, as described below. Total LRRK2 and Rab10 levels were determined as a reference.

For Western Blot analysis, protein samples were separated by SDS–PAGE using NuPAGE 10% Bis-Tris gels (Invitrogen) and transferred onto PVDF membranes (Thermo Fisher).

To allow simultaneous probing for LRRK2 on the one hand and Rab and the Nb-GFP fusions on the other hand, membranes were cut horizontally at the 55 and 140 kDa MW marker band. After blocking non-specific binding sites with 5% non-fat dry milk in TBS-T (1 h, RT) (25 mM Tris, pH 7.4, 150 mM NaCl, 0.1% Tween-20), membranes were incubated overnight at 4°C with primary antibodies at dilutions specified below. Phospho-specific antibodies were diluted in TBST/ 5% BSA (Roth GmbH). Non-phospho-specific antibodies were diluted in TBST/ 5% non-fat dry milk powder (CST). Phospho-Rab10 levels were determined by the site-specific rabbit monoclonal antibody anti-pRAB10(pT73) (Abcam, ab230261) and LRRK2 autophosphorylation was determined by the site-specific rabbit monoclonal antibody anti-pLRRK2(pS1292) (Abcam, ab203181), both at a dilution of 1:2,000. Total LRRK2 levels were determined by an in-house rat monoclonal antibody (clone 24D8; 1:5000) (11). Total Rab10 levels were determined by the rabbit monoclonal antibody anti-RAB10/ERP13424 (Abcam, ab181367) at a dilution of 1:5,000. Nb-GFP fusion proteins were detected using the rat monoclonal antibody anti-GFP (clone 3H9, ChromoTec) at a dilution of 1:2,000. For detection, goat anti-rat IgG or anti-rabbit IgG HRP-coupled secondary antibodies (Jackson ImmunoResearch) were used at a dilution of 1:15,000 in TBST/ 5% non-fat dry milk powder. Antibody–antigen complexes were visualized using the ECL plus chemiluminescence detection system (GE Healthcare) on Hyperfilms (GE Healthcare).

**Chemical crosslinking/ mass spectrometry (CL-MS)**

The chemical crosslinking of the LRRK2 protein with each of the Nbs, using the NHS-ester-based and CID-cleavable reagent disuccinimidyl sulfoxide (DSSO; Thermo Fisher) (12), was performed as outlined in the Materials and Methods section of the main text. After crosslinking, precipitation, tryptic proteolysis, clean-up and enrichment for crosslinked peptides (1), vacuum-dried fractions containing the crosslinked peptides, were analyzed individually on an Orbitrap Fusion mass spectrometer (Thermo Fisher) using the MS2_MS3 fragmentation method with the default settings (ver. 3.0, build 2041). MS1 scans were performed in the Orbitrap (FTMS, resolution = 60K) at an m/z range of 375-1500. MS2 was performed with CID (CE=25%) and spectra were acquired in the Orbitap (FTMS) at 30K resolution. The MS3 scans were performed with HCD (CE=30%) and spectra were acquired in the linear ion trap. The Thermo Raw files were analyzed with the

MS2_MS3 workflow provided by in Proteome Discoverer 2.5 (build 2.5.0.400), which uses XlinkX (ver. 2.5) (13) for the detection of crosslinked peptides. A global search of MS2 spectra was performed with Sequest HT against the human subset of the Swissprot database (v. 2019_02; 20417 entries) supplemented with the sequences of the nanobodies followed by an FDR analysis (FDR=0.01) by the Target Decoy PSM validator. For the Sequest analysis, the following settings have been used: Trypsin has been used as enzyme. Carbamylation of cysteines has been used as fixed modification and Methionine oxidation, DSSO hydrolyzed (K+176.014 Da), DSSO Tris (K+279.078 Da) and n-terminal Acetylation were allowed as variable modifications.

For the detection of crosslinked peptides by the XlinkX detection node, the acquisition strategy was set to MS2_MS3 and DSSO (158.004; K) was used as crosslinker at an S/N minimum of 1.5. For the XlinkX database search, the following parameters have been used: Trypsin has been used as enzyme. The precursor and fragment mass tolerances were set to 10 ppm (precursor), 20 ppm (FTMS) and 0.5 Da (ITMS), respectively. The search was performed using a database containing the LRRK2 sequence and the individual sequences of all used nanobodies, including an irrelevant control (total: 11 sequences). Carbamidometyl has been used as fixed and Methionine oxidation was allowed as variable modification. FDR-based analysis (XlinkX/PD validator node) was performed using an FDR threshold of 0.01. In addition, only XlinkX scores>15 were considered.

In the consensus step, the identified crosslinks were filtered for an identification score ≥20 (default value) to reduce the number of false-positive hits. Filtered crosslinking data were exported and visualized in xiNet (14).

### Supplementary Figures & Tables

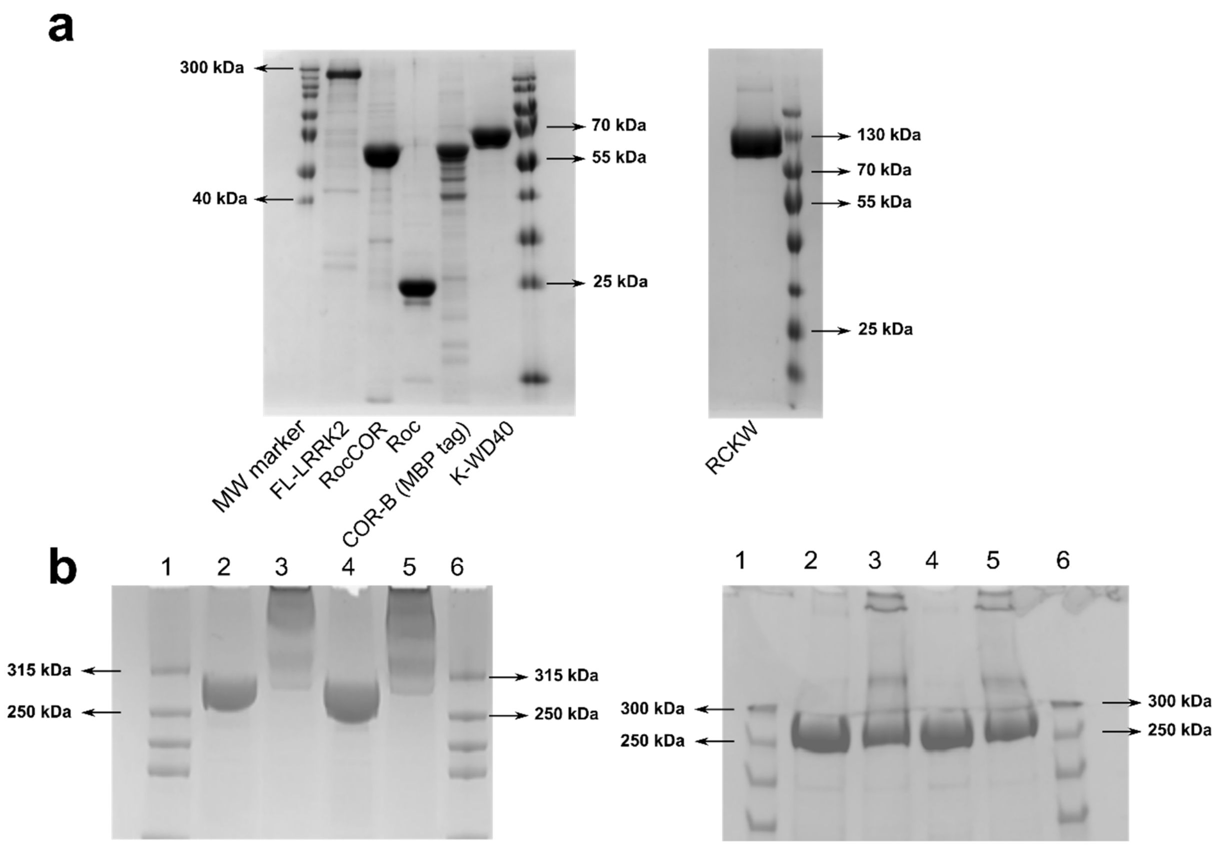


**Fig. S1. Purification and crosslinking of LRRK2 and LRRK2 constructs. (a)** SDS-PAGE analysis of purified full-length LRRK2 and LRRK2 domain constructs used in this study. 3 µg (5 µg for RCKW) of each purified protein, FL-LRRK2, RocCOR, Roc, (MBP-)COR-B, kinase-WD40 and RCKW, was loaded on gel. (**b**) Crosslinking of LRRK2 with the lysine-specific crosslinker DSS, as used for immunization. Two cross-link setups were performed, where cross-linking was allowed to proceed to different levels (*left* and *right* panels). LRRK2 was purified in presence of GDP (lanes 2 and 3) or GTPγS (lanes 4 and 5). Lanes 2 and 4 show LRRK2 before cross-linking, while lanes 3 and 5 show LRRK2 after cross linking. For immunization 2, a mixture of the samples shown in lanes 5 was used, for immunization 3 a mixture of the samples shown in lanes 3 was used.

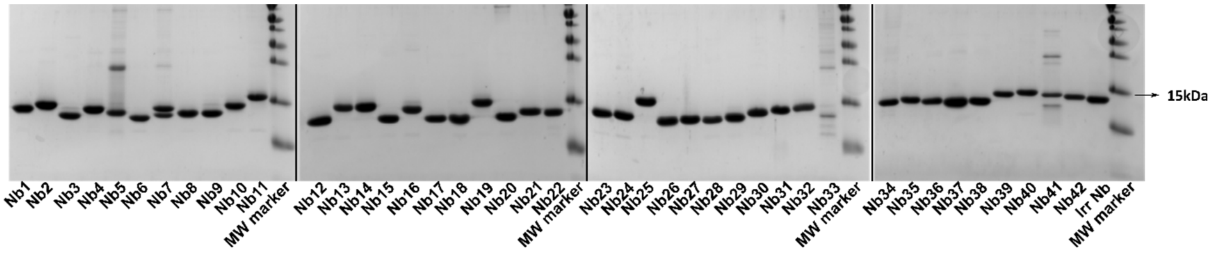


**Fig. S2. SDS-PAGE of the 42 purified Nbs and the irrelevant Nb used in this study.** 2 µg of each Nb was loaded on gel (except for Nb33 where 0.7 µg is loaded).

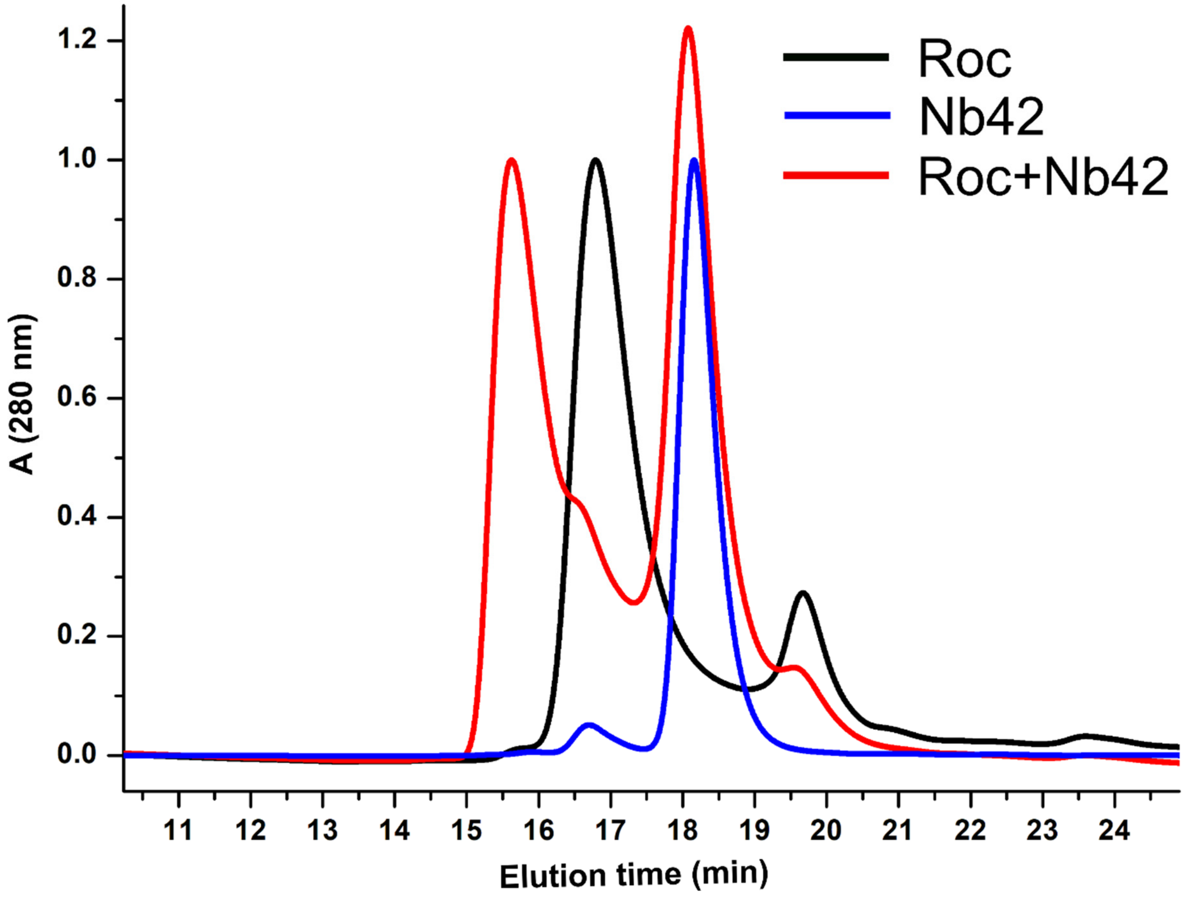


**Fig. S3.** Size exclusion chromatogram of the LRRK2 Roc domain (black curve), Nb42 (blue curve), and the Roc domain mixed with Nb42 (red curve), showing a shift to lower elution time and higher molecular weight upon binding of the Nb to the Roc domain. The LRRK2 Roc domain (residues 1329-1520) was diluted to a concentration of 50 µM and either mixed or not with a molar excess of Nb42. After 30 min incubation both samples were injected on a Bio-SEC 3 size exclusion chromatography column (Agilent) connected to an HPLC system (Waters), using 30 mM HEPES pH 7.5, 150 mM NaCl, 10 mM $MgCl_2$, 5% glycerol and 100 mM GDP as a running buffer.

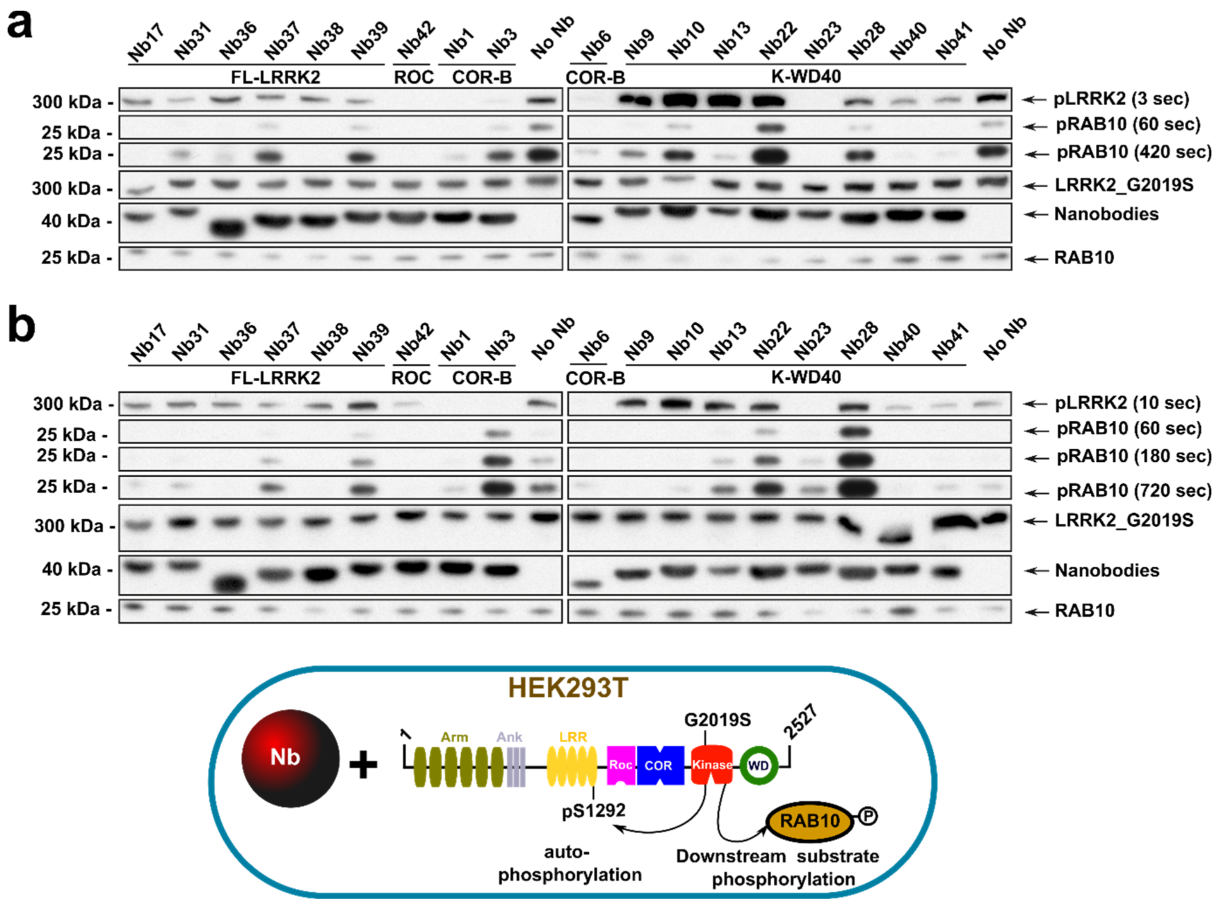


**Fig. S4. Influence of Nbs on the kinase activity of the LRRK2(G2019S) variant in HEK293T cells** (additional repeats of experiment shown in Fig. 2b). **(a-b)** LRRK2(G2019S) and its effector Rab29 were overexpressed together with GFP-tagged Nbs in HEK293T cells. A negative control, where no Nb is overexpressed ("No Nb") is also included. Panel (a) and (b) show two additional repeats of the experiments shown in Fig. 2b. In rows labelled "pLRRK2", LRRK2 pS1292 levels are determined by Western Blot using a site-specific anti-pLRRK2(pS1292) antibody (Abcam, ab203181) (shown at different times of development). In the rows labeled "pRAB10", endogenous pT73-Rab10 levels are determined by Western Blot using the MJFF/Abcam antibody MJF-R21 (Abcam, ab230261) (shown at different times of development). The three lower rows contain controls of LRRK2 (rat anti-LRRK2, 24D8), GFP-Nb (rat anti-GFP) and Rab10 (rabbit anti-Rab10) expression levels, determined on a different Western blot than pLRRK2 and pRab10.

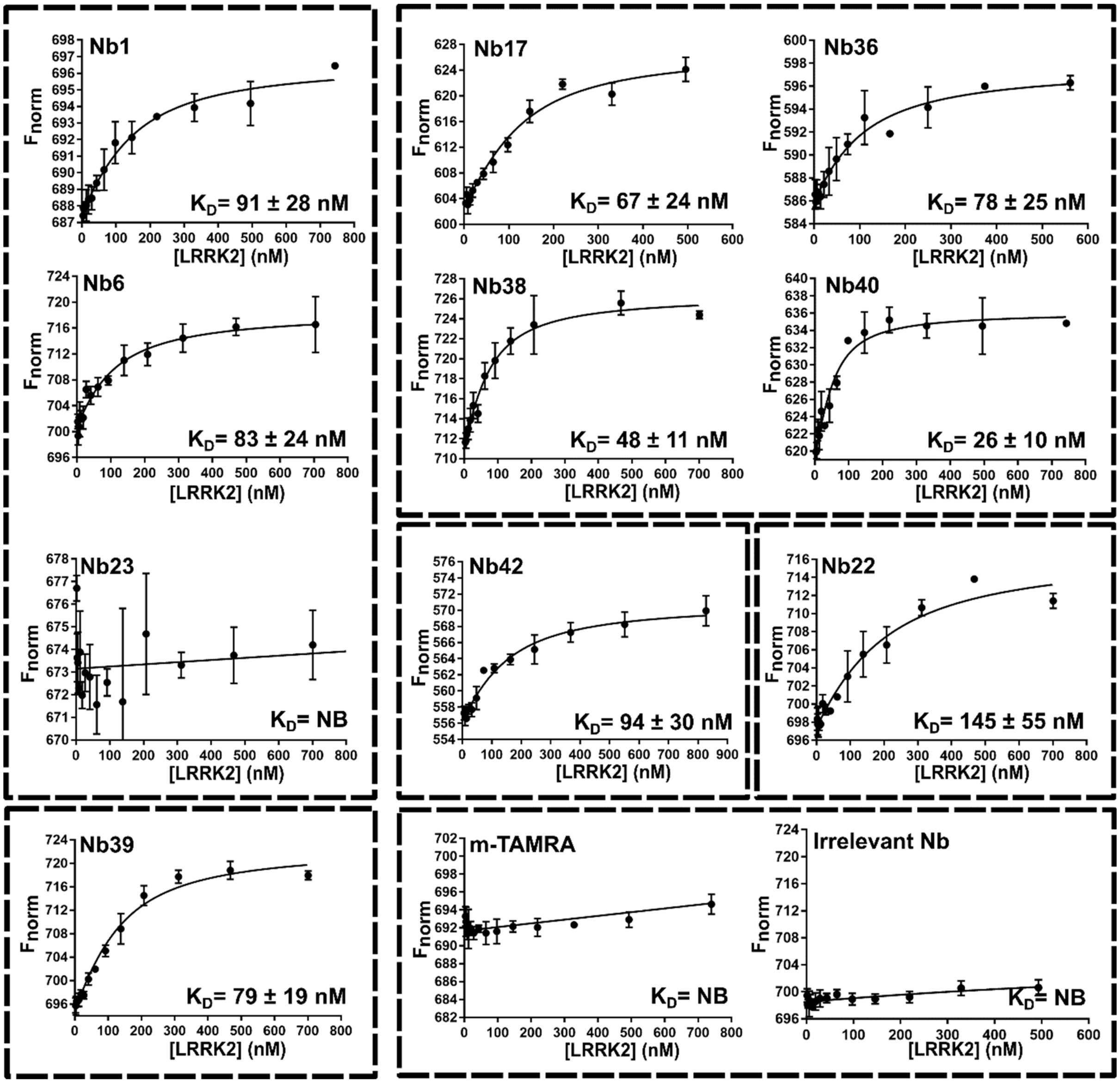


**Fig. S5. Affinity measurements of the Nbs for LRRK2 using Microscale thermophoresis (MST).** Binding isotherms are shown, obtained by titrating increasing concentrations of LRRK2 to fluorescently (m-TAMRA)- labeled Nbs. Nbs are classified into five functional groups, as defined in Fig. 1b with group1 Nbs: Nb1, Nb6, Nb23; group2 Nbs: Nb17, Nb36, Nb38 and potentially Nb40; group3 Nb: Nb42; group4 Nb: Nb22; and group5 Nbs: Nb39 and potentially Nb40. In the lower right panel, two negative controls are shown, where LRRK2 was either titrated to free m-TAMRA or to a m-TAMRA-labeled irrelevant Nb. All measurements were performed in presence of 500 μM GDP, except for Nb42 where GTPγS was used. The $K_D$ values (± standard error) from fitting on a quadratic binding equation are given (each data point is the average ± SD of n=3; NB = no MST binding signal detectable).

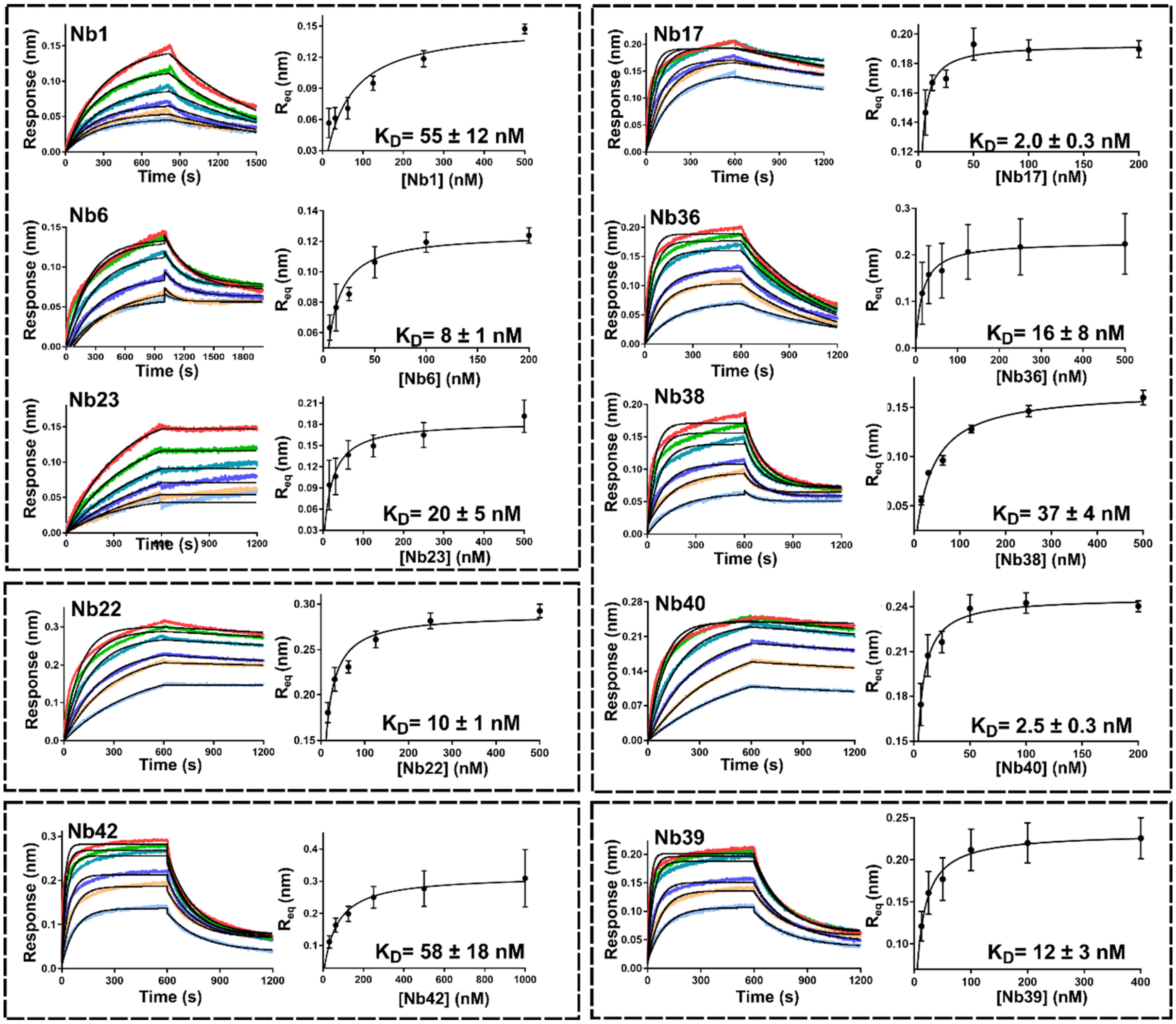


**Fig. S6. Affinity measurements of the Nbs for LRRK2 using Biolayer Interferometry (BLI).** BLI sensorgrams obtained by titrating increasing concentrations of the Nbs to LRRK2 that was trapped on a Streptavidine biosensor via biotinylated Nb40 (or Nb42 in case of affinity measurement of Nb40) (*left*), and the resulting dose-response curves (*right*) are shown. The sensorgrams were fitted on a 1:1 binding model (ForteBio Analysis Software) and the resulting $R_{eq}$ values were subsequently plotted against the Nb concentration. The $K_D$ values (± standard error) obtained by fitting the dose-response curves with a Langmuir binding equation are given (each data point is the average ± SD of n=3). Nbs are classified into five functional groups as defined in Fig. 1b with group1 Nbs: Nb1, Nb6, Nb23; group2 Nbs: Nb17, Nb36, Nb38 and potentially Nb40; group3 Nb: Nb42; group4: Nb: Nb22; and group5 Nb: Nb39 and potentially Nb40. All measurements were performed in presence of 500 µM GDP.

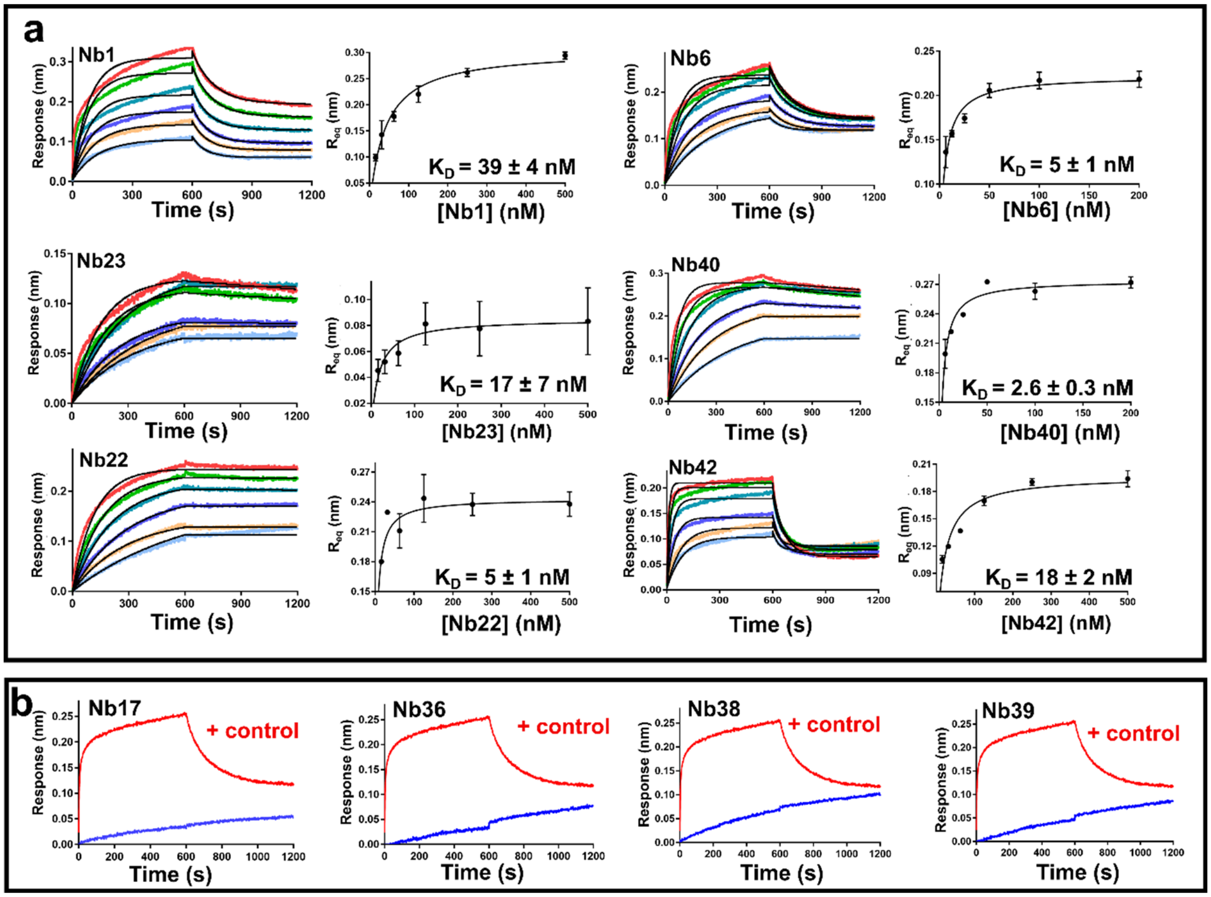


**Fig. S7. Affinity measurements of the Nbs for the LRRK2 RCKW construct using Biolayer Interferometry (BLI). (a)** BLI sensorgrams obtained by titrating increasing concentrations of the Nbs to biotinylated RCKW that was trapped on a Streptavidine biosensor (*left*), and the resulting dose-response curves (*right*) are shown. All measurements were performed in presence of 500 µM GDP. Analysis was performed as in Fig. S6. The $K_D$ values (± standard error) obtained by fitting with a Langmuir binding equation are given (each data point is the average ± SD of n=3). **(b)** Nb17, Nb36, Nb38 and Nb39 that were identified as "full-length LRRK2 binders" in ELISA and CL-MS correspondingly do not bind to the RCKW construct (blue curves). A positive control experiment with Nb6 was performed and shown as the red curve.

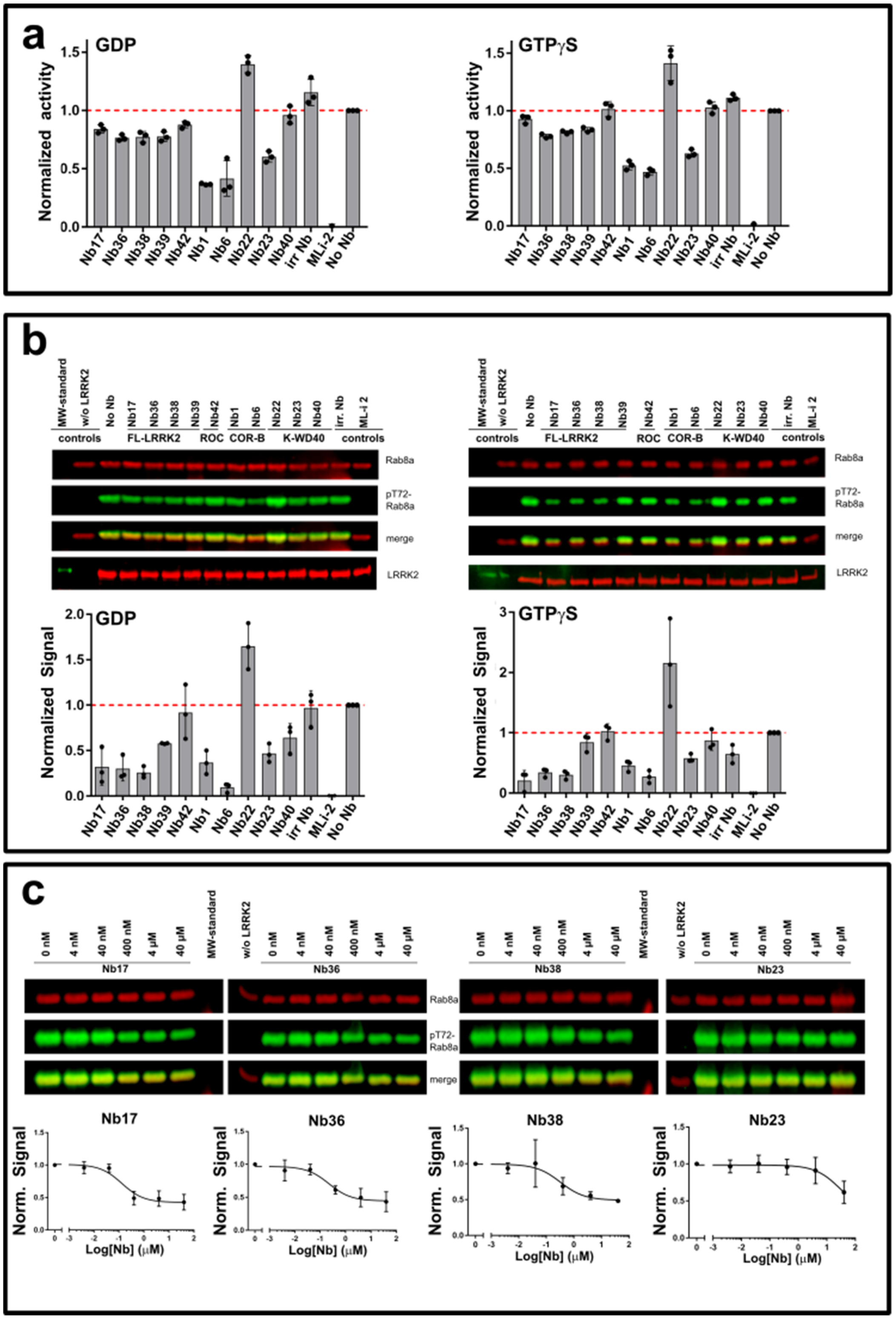


**Fig. S8. Modulation of *in vitro* LRRK2 kinase activity by the Nbs. (a)** Effect of Nbs on LRRK2 kinase activity measured using the LRRK2-optimized AQT0615 peptide (at 10 µM) as substrate, either in the presence of 500 µM GDP (*left*) or GTPgS (*right*). (b) Effect of Nbs on LRRK2-

mediated phosphorylation of Rab8a (at 2.5 μM) determined via a Western blot assay, either in the presence of 500 μM GDP (*left*) or GTPgS (*right*). The ratio of the pT72-Rab8a signal to the total Rab8a signal was used for quantification. In both (a) and (b) the influence of the Nbs (25 μM) on the relative kinase activity compared to the "No-Nb" control is plotted, and a positive control with 0.25 μM MLi-2 is included. Each bar reflects the average (± SD) of three independent measurements. (c) Semi-quantitative dose-response curves for Nb17, Nb36 Nb38 and Nb23 (*lower panel*) derived from the respective normalized signal of Rab8a phosphorylation (*upper panel*) performed with varying Nb concentration from 4 nM to 40 μM (10-fold serial dilution). The dose-response curves represent the average of four (± SD) independent measurements.

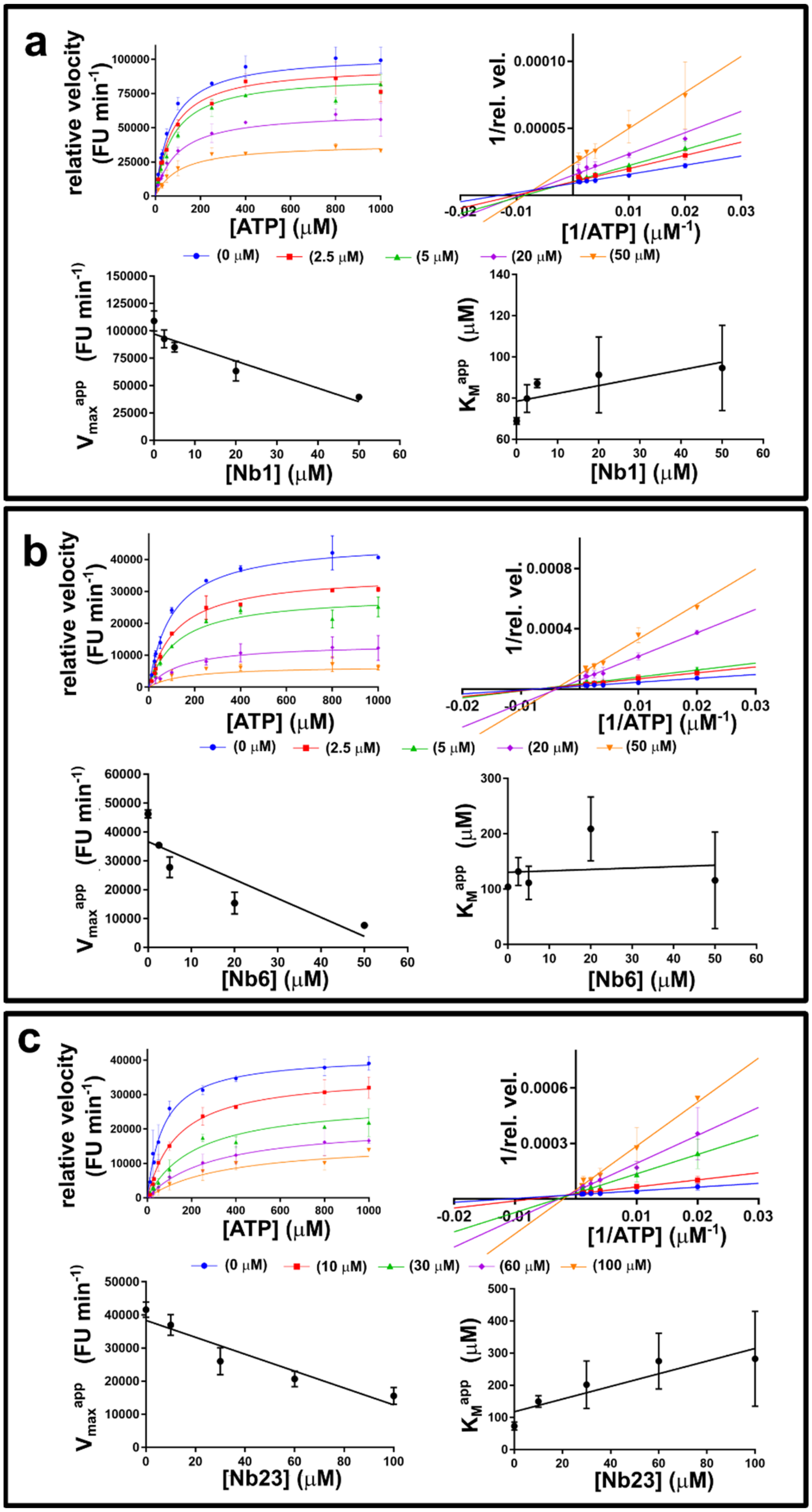


**Fig. S9. Mechanism of inhibition of the group1 Nbs: Nb1, Nb6, Nb23.** Global fitting and analysis of Michaelis-Menten curves for LRRK2 kinase activity obtained at varying concentrations of ATP, a fixed (sub-saturating) concentration of peptide substrate (AQT0615) and at varying concentrations of Nb1 (**a**), Nb6 (**b**) and Nb23 (**c**). The upper figures of each panel show the results of the global fit on a mixed-inhibition model and the linearization of the Michaelis-Menten curves according to the Lineweaver-Burk method (double-reciprocal plot). The lower panels show the influence of the Nb concentration of the apparent maximal velocities ($V_{max}^{app}$) and apparent $K_M$ values ($K_M^{app}$), corresponding to each individual velocity *versus* [Nb] trace. Corresponding to a mixed-type inhibition model $V_{max}^{app}$ decreases and $K_M^{app}$ increases (or remains constant) with increasing Nb concentration. The Nb concentrations used are plotted in different color and the values are indicated. Each datapoint reflects the average (± standard error) of three independent measurements.

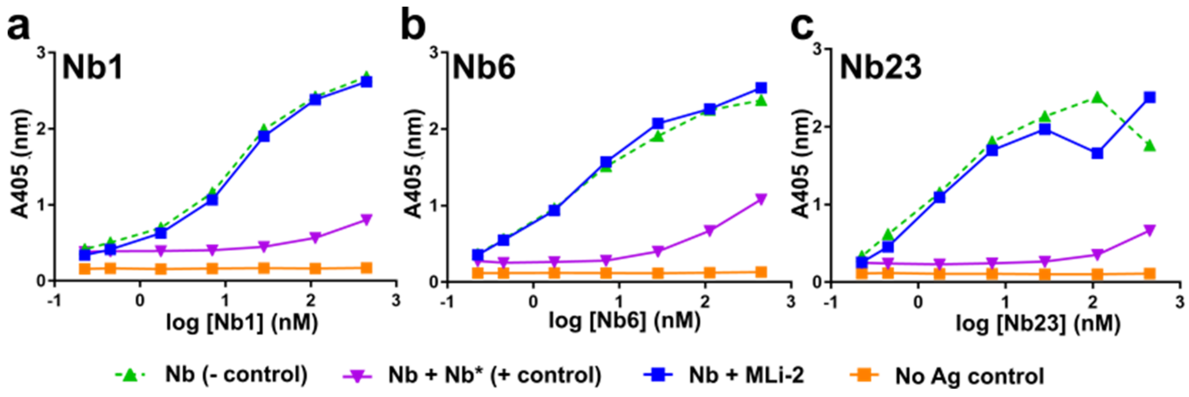


**Fig. S10. Nbs bind LRRK2 through binding sites differing from the ATP/MLi-2 binding pocket.** Competition ELISA titration experiments assessing whether the ATP competitive kinase inhibitor MLi-2 competes for the same binding sites as the group1 Nbs: Nb1 **(a),** Nb6 **(b)** and Nb23 **(c).** LRRK2 was coated in the wells of the ELISA plate and the ELISA signal for a dilution series of the respective Nbs (detected via their C-terminal EPEA-tag) is plotted in function of the Nb concentration. The effect of the presence of a large excess of MLi-2 (1 µM) and the corresponding untagged Nb as a positive (+) control (Nb*, at 9 µM) is determined. A "no antigen control", where no LRRK2 was coated on the bottom of the well, is also included. In contrast to the positive control where addition of Nb* causes a clear rightward shift of the ELISA titration curves compared to the titration curve of the Nb alone ("- control"), MLi-2 does not show a rightward shift of the curves.

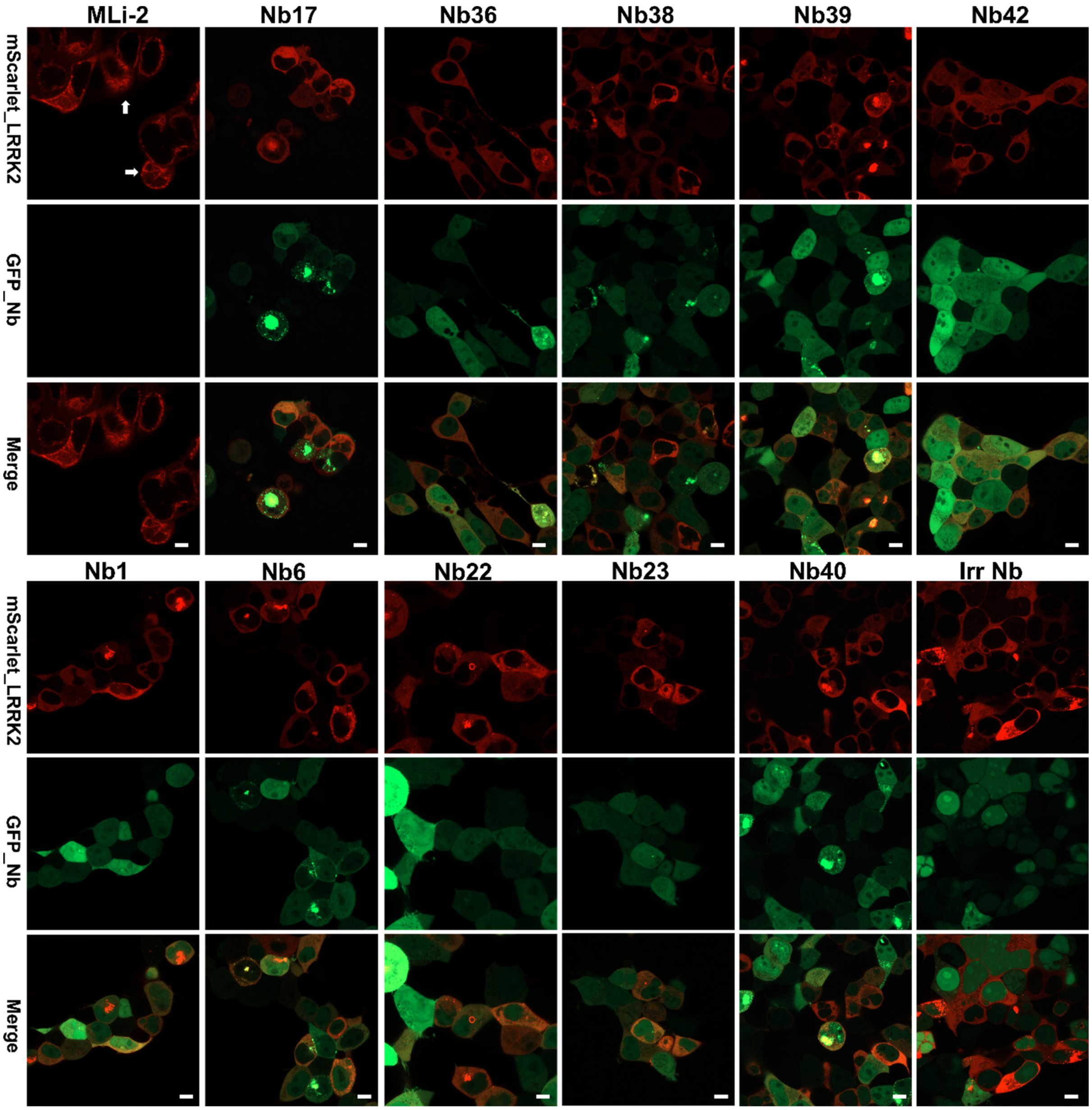


**Fig. S11. Effect of LRRK2-targeting Nbs on cellular localization of LRRK2.** HEK293 cells were co-transfected with the indicated GFP-Nbs and mScarlet-LRRK2. As a positive control mScarlet-LRRK2-transfected cells were treated with the pharmacological ATP-competitive inhibitor MLi-2 (1 µM MLi-2, 90 mins treatment), showing induction of LRRK2 relocalization onto microtubules seen as filamentous skein-like structures and indicated by white arrows (upper left panel). In contrast, cells co-transfected with GFP-Nbs and mScarlet-LRRK2 showed normal cytoplasmic distribution of LRRK2 with no relocalization to microtubules, similar to cells co-transfected with an irrelevant Nb. Scale bar 5 µm.

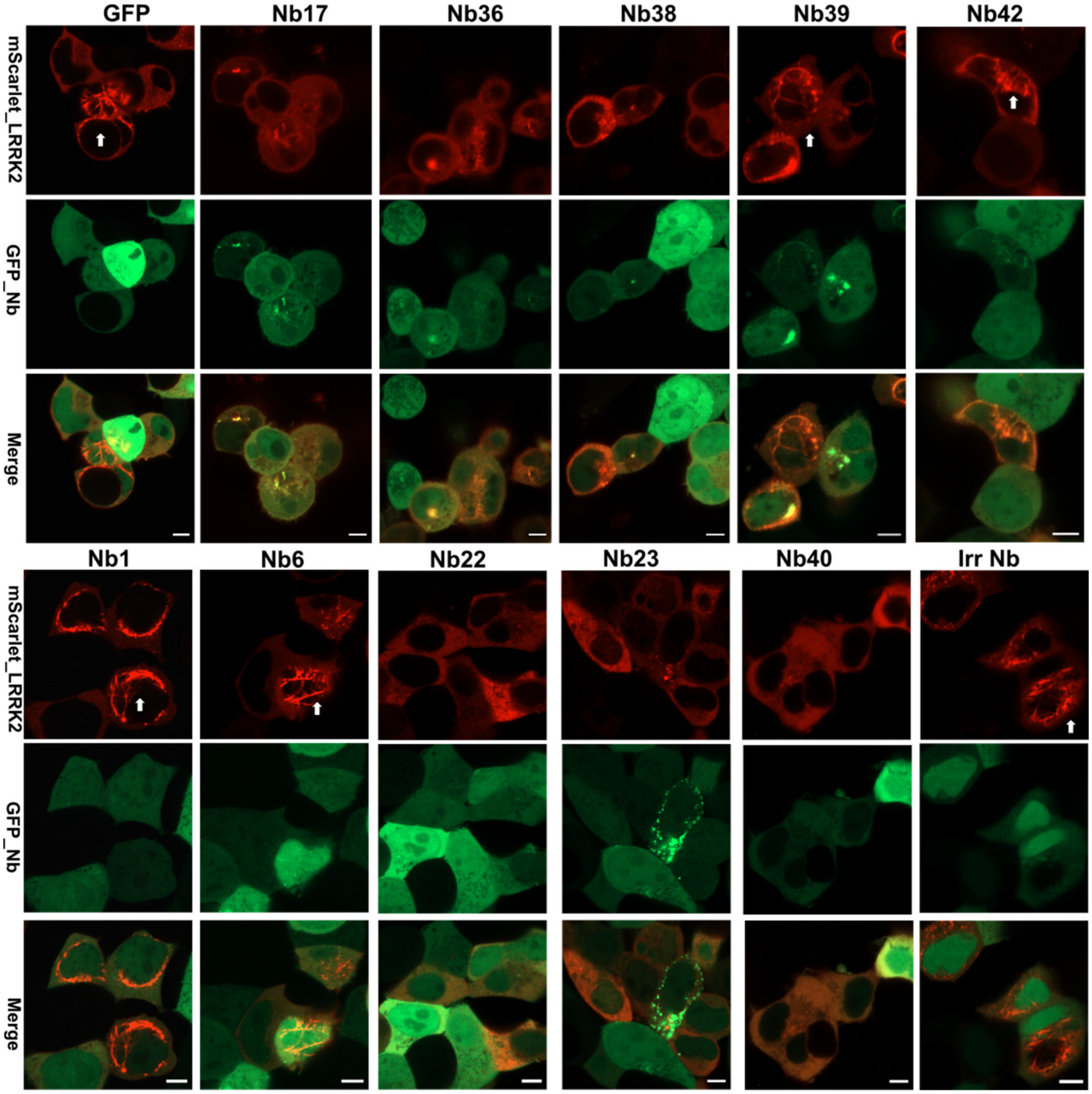


**Fig. S12. Effect of LRRK2-targeting Nanobodies on MLi-2-induced microtubule relocalization of LRRK2.** HEK293 cells were co-transfected with the indicated GFP-Nbs and mScarlet-LRRK2 and treated with the pharmacological ATP-competitive inhibitor MLi-2 (1 µM MLi-2, 90 mins treatment). Co-transfection of mScarlet-LRRK2 with only GFP (upper left panel) or an irrelevant Nb (lower left panel) showed the MLi-2-induced relocalization of LRRK2 onto microtubules seen as filamentous skein-like structures and indicated by white arrows. Co-transfection with a subset of GFP-Nb inhibited the MLi-2-induced LRRK2 relocalization. Scale bar 5 µm.

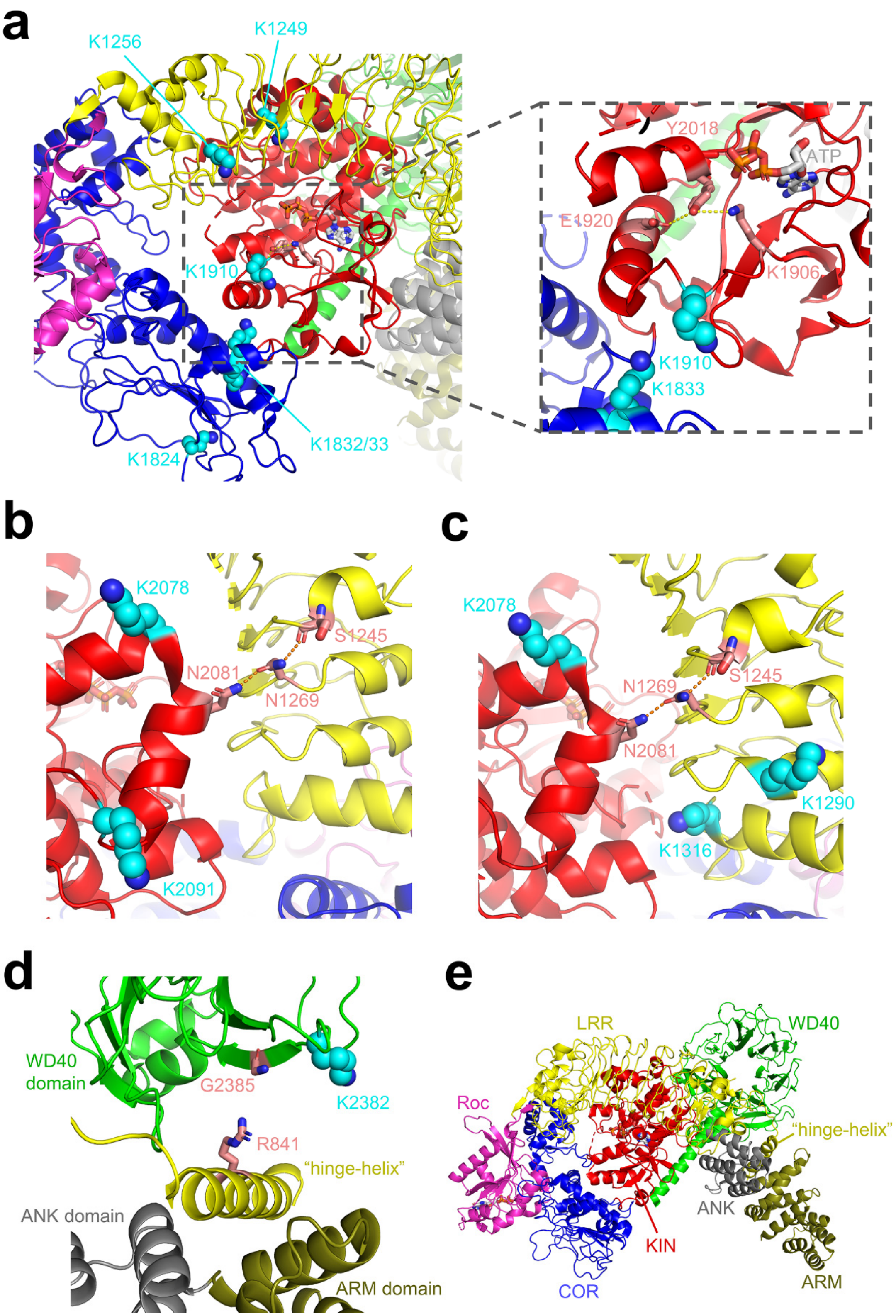


**Fig. S13. Zoomed in representations of the cross-linking sites of Nbs belonging to group 1 (Nb1 and Nb23), group 2 (Nb17) and group 3 (Nb22)** (see main document Fig. 6 for an overall view of the other Nb representatives). In panels (a) – (d) the lysine residues of LRRK2 that make cross-links to the respective Nbs are shown as spheres with carbon atoms colored cyan, and other relevant residues discussed in the main text are shown as sticks with carbon atoms colored light red. The different LRRK2 domains are colored as indicated in panel (e). **(a)** Cross-linking pattern of Nb1, belonging to the inhibitory Nb group 1. While ELISA experiments showed that this Nb is able to bind exclusively to the C-terminal part of the COR domain (COR-B), cross-links are made with residues from the LRR, COR and kinase domains, suggesting a binding position in a central cavity of the LRRK2 structure. The zoom-in figure on the right shows that the cross-linking sites K1833 and K1910 are located in the vicinity of K1906 and E1920 and could potentially stabilize the inhibitory K1906-Y2018-E1920 hydrogen bonding network. **(b)** Cross-linking pattern of Nb23, belonging to the inhibitory Nb group 1. The cross-linking sites K2078 and K2091 surround residue N2081. The kinase activating N2081D mutation was identified as a risk factor for Crohn's disease (15), potentially by disrupting the N2081-N1269 hydrogen bond which contributes to keeping the LRR domain in an inhibitory conformation. Likewise, one could speculate that Nb23 inhibits kinase activity by stabilizing this inhibitory LRR conformation. **(c)** Cross-linking pattern of Nb17, belonging to the inhibitory Nb group 2. Nb17 binds on the interface of several LRRK2 domains ("FL-LRRK2 binder") and cross-link with LRR residues K1290 and K1316 and kinase domain residue K2078, again in proximity of N2081. Similar to Nb23, Nb17 could stabilize the LRR domain in a conformation that inhibits binding and phosphorylation of bulky substrates, while in contrast to Nb23, still allowing auto-phosphorylation and phosphorylation of small peptide substrates. **(d)** Cross-linking pattern of Nb22, belonging to the kinase-activating Nb group 3. Nb22 cross-links with K2382 within the WD40 domain, very close to the site of the G2385R PD risk variant (16). The latter mutation activates kinase activity presumably by disrupting the (inhibitory) interaction with the LRR "hinge helix". One could speculate that Nb22 has the same mechanism of kinase activation as G2385R. **(e)** Overall view on the cryo-EM structure of full-length LRRK2 (PDB 7lhw, (17) illustrating the color codes that were used to indicate the different LRRK2 domains: armadillo domain: dark yellow, ankyrin domain: grey, leucin-rich repeat domain: yellow, Roc domain: magenta, COR domain blue, kinase domain: red, WD40 domain: green. The LRR "hinge-helix" is also indicated. ATP (bound to the kinase domain) and GDP (bound to the Roc domain) are shown in ball-and-stick representations with carbon atoms colored grey.

**Table S1:** List of purified Nbs resulting from: (1) the immunization with RocCOR in presence of GppNHp (Immunization 1) and selection with full-length LRRK2, (2) the immunization with LRRK2-GTPγS (Immunization 2) and selection with either LRRK2-GTPγS or Roc-GTPγS, and (3) the immunization with LRRK2-GDP (Immunization 3) and selection with either LRRK2-GDP or Roc-GDP. The domain specificity determined using ELISA is indicated. Nbs investigated in this study are indicated in bold.

| Nb | Internal database nr. | Immunization | Selection | Target Domain ELISA |
|---|---|---|---|---|
| **Nb1** | **CA12610** | **Imm. 1 - RocCOR-GppNHp** | **LRRK2-GDP** | **COR-B** |
| Nb2 | CA12612 | Imm. 1 - RocCOR-GppNHp | LRRK2-GDP | COR-B |
| **Nb3** | **CA12614** | **Imm. 1 - RocCOR-GppNHp** | **LRRK2-GDP** | **COR-B** |
| Nb4 | CA12616 | Imm. 1 - RocCOR-GppNHp | LRRK2-GDP | COR-B |
| Nb5 | CA12617 | Imm. 1- RocCOR-GppNHp | LRRK2-GDP | COR-B |
| **Nb6** | **CA12618** | **Imm. 1 - RocCOR-GppNHp** | **LRRK2-GDP** | **COR-B** |
| Nb7 | CA12620 | Imm. 1 - RocCOR-GppNHp | LRRK2-GDP | COR-B |
| Nb8 | CA13597 | Imm. 2 - LRRK2-GTPγS | LRRK2-GTPγS | K-WD40 |
| **Nb9** | **CA13598** | **Imm. 2 - LRRK2-GTPγS** | **LRRK2-GTPγS** | **K-WD40** |
| **Nb10** | **CA13599** | **Imm. 2 - LRRK2-GTPγS** | **LRRK2-GTPγS** | **K-WD40** |
| Nb11 | CA13600 | Imm. 2 - LRRK2-GTPγS | LRRK2-GTPγS | K-WD40 |
| Nb12 | CA13601 | Imm. 2 - LRRK2-GTPγS | LRRK2-GTPγS | K-WD40 |
| **Nb13** | **CA13602** | **Imm. 2 - LRRK2-GTPγS** | **LRRK2-GTPγS** | **K-WD40** |
| Nb14 | CA13603 | Imm. 2 - LRRK2-GTPγS | LRRK2-GTPγS | K-WD40 |
| Nb15 | CA13604 | Imm. 2 - LRRK2-GTPγS | LRRK2-GTPγS | K-WD40 |
| Nb16 | CA13605 | Imm. 2 - LRRK2-GTPγS | LRRK2-GTPγS | K-WD40 |
| **Nb17** | **CA13606** | **Imm. 2 - LRRK2-GTPγS** | **LRRK2-GTPγS** | **FL-LRRK2** |
| Nb18 | CA13607 | Imm. 2 - LRRK2-GTPγS | LRRK2-GTPγS | FL LRRK2 |
| Nb19 | CA13608 | Imm. 2 - LRRK2-GTPγS | LRRK2-GTPγS | K-WD40 |
| Nb20 | CA13609 | Imm. 2 - LRRK2-GTPγS | LRRK2-GTPγS | K-WD40 |
| Nb21 | CA13610 | Imm. 2 - LRRK2-GTPγS | LRRK2-GTPγS | K-WD40 |
| **Nb22** | **CA13611** | **Imm. 2 - LRRK2-GTPγS** | **LRRK2-GTPγS** | **K-WD40** |
| **Nb23** | **CA13612** | **Imm. 2 - LRRK2-GTPγS** | **LRRK2-GTPγS** | **K-WD40** |
| Nb24 | CA13613 | Imm. 2 - LRRK2-GTPγS | LRRK2-GTPγS | FL-LRRK2 |
| Nb25 | CA13614 | Imm. 2 - LRRK2-GTPγS | LRRK2-GTPγS | No binding |
| Nb26 | CA13615 | Imm. 2 - LRRK2-GTPγS | LRRK2-GTPγS | FL-LRRK2 |
| Nb27 | CA13616 | Imm. 2 - LRRK2-GTPγS | LRRK2-GTPγS | K-WD40 |
| **Nb28** | **CA13617** | **Imm. 2 - LRRK2-GTPγS** | **LRRK2-GTPγS** | **K-WD40** |
| Nb29 | CA13618 | Imm. 2 - LRRK2-GTPγS | LRRK2-GTPγS | No binding |
| Nb30 | CA13619 | Imm. 2 - LRRK2-GTPγS | LRRK2-GTPγS | FL-LRRK2 |
| **Nb31** | **CA13620** | **Imm. 2 - LRRK2-GTPγS** | **LRRK2-GTPγS** | **FL-LRRK2** |
| Nb32 | CA16069 | Imm. 2 - LRRK2-GTPγS | Roc-GTPγS | Roc |
| Nb33 | CA16070 | Imm. 2 - LRRK2-GTPγS | Roc-GTPγS | Roc |
| Nb34 | CA16071 | Imm. 2 - LRRK2-GTPγS | Roc-GTPγS | Roc |
| Nb35 | CA16072 | Imm. 2 - LRRK2-GTPγS | Roc-GTPγS | Roc |
| **Nb36*** | **CA14130** | **Imm. 3 - LRRK2-GDP** | **LRRK2-GDP** | **FL-LRRK2** |
| **Nb37** | **CA14131** | **Imm. 3 - LRRK2-GDP** | **LRRK2-GDP** | **FL-LRRK2** |
| **Nb38*** | **CA14133** | **Imm. 3 - LRRK2-GDP** | **LRRK2-GDP** | **FL-LRRK2** |
| **Nb39** | **CA14134** | **Imm. 3 - LRRK2-GDP** | **LRRK2-GDP** | **FL-LRRK2** |
| **Nb40** | **CA14135** | **Imm. 3 - LRRK2-GDP** | **LRRK2-GDP** | **K-WD40** |
| **Nb41** | **CA14136** | **Imm. 3 - LRRK2-GDP** | **LRRK2-GDP** | **K-WD40** |
| **Nb42** | **CA14259** | **Imm. 3 - LRRK2-GDP** | **Roc-GDP** | **Roc** |

* Nb36 and Nb38 belong to the same sequence family with only a number of amino acid substitutions in the CDR regions

### SI References


1. G. Guaitoli, *et al.*, Structural model of the dimeric Parkinson's protein LRRK2 reveals a compact architecture involving distant interdomain contacts. *Proc. Natl. Acad. Sci.* **113**, E4357–E4366 (2016).
2. C. J. Gloeckner, *et al.*, Phosphopeptide analysis reveals two discrete clusters of phosphorylation in the N-terminus and the Roc domain of the Parkinson-disease associated protein kinase LRRK2. *J. Proteome Res.* **9**, 1738–45 (2010).
3. C. J. Gloeckner, K. Boldt, A. Schumacher, R. Roepman, M. Ueffing, A novel tandem affinity purification strategy for the efficient isolation and characterisation of native protein complexes. *Proteomics* **7**, 4228–34 (2007).
4. N. Gul, D. M. Linares, F. Y. Ho, B. Poolman, Evolved escherichia coli strains for amplified, functional expression of membrane proteins. *J. Mol. Biol.* **426**, 136–49 (2014).
5. F. Y. Ho, B. Poolman, Engineering escherichia coli for functional expression of membrane proteins. *Methods Enzymol.* **556**, 3–21 (2015).
6. E. Pardon, *et al.*, A general protocol for the generation of Nanobodies for structural biology. *Nat. Protoc.* **9**, 674–693 (2014).
7. H. L. Ulrich Rothbauer, Kourosh Zolghadr, Sergei Tillib, Danny Nowak, Lothar Schermelleh, Anja Gahl, Natalija Backmann, Katja Conrath, Serge Muyldermans, M Cristina Cardoso, Targeting and tracing antigens in live cells with fluorescent nanobodies. *Nat. Methods* **3**, 887–889 (2006).
8. B. Traenkle, *et al.*, Monitoring interactions and dynamics of endogenous beta-catenin with intracellular nanobodies in living cells. *Mol. Cell. Proteomics* **14**, 707–723 (2015).
9. J. Maier, B. Traenkle, U. Rothbauer, Real-time analysis of epithelial-mesenchymal transition using fluorescent single-domain antibodies. *Sci. Rep.* **5** (2015).
10. C. J. Gloeckner, K. Boldt, M. Ueffing, Strep/FLAG tandem affinity purification (SF-TAP) to study protein interactions. *Curr. Protoc. Protein Sci.* **57**, 19.20.1-19.20.19 (2009).
11. M. D. P. Carrion, *et al.*, The LRRK2 G2385R variant is a partial loss-of-function mutation that affects synaptic vesicle trafficking through altered protein interactions. *Sci. Rep.* **7**, 5377 (2017).
12. W. V. Kandur, A. Kao, D. Vellucci, L. Huang, S. D. Rychnovsky, Design of CID-cleavable protein cross-linkers: Identical mass modifications for simpler sequence analysis. *Org. Biomol. Chem.* **13**, 9793–807 (2015).
13. F. Liu, D. T. S. Rijkers, H. Post, A. J. R. Heck, Proteome-wide profiling of protein assemblies by cross-linking mass spectrometry. *Nat. Methods* **12**, 1179–84 (2015).
14. C. W. Combe, L. Fischer, J. Rappsilber, xiNET: Cross-link network maps with residue resolution. *Mol. Cell. Proteomics* **14**, 1137–47 (2015).
15. K. Y. Hui, *et al.*, Functional variants in the LRRK2 gene confer shared effects on risk for Crohn's disease and Parkinson's disease. *Sci.Transl.Med.* **10**, eaai7795 (2018).
16. I. F. Mata, *et al.*, Lrrk2 pathogenic substitutions in Parkinson's disease. *Neurogenetics* **6**, 171–177 (2005).
17. A. Myasnikov, *et al.*, Structural analysis of the full-length human LRRK2. *Cell* **184**, 1–9 (2021).
